# Scoring information integration with statistical quality control enhanced cross-run analysis of data-independent acquisition proteomics data

**DOI:** 10.1101/2024.12.19.629475

**Authors:** Mingxuan Gao, Shubham Gupta, Wenxian Yang, Rongshan Yu, Hannes L. Röst

**Author notes:** DIA proteomics | cross-run | match-between-runs | deep learning | signal alignment.

## Abstract

The peptide-centric strategy is widely applied in data-independent acquisition (DIA) proteomics to analyze multiplexed MS2 spectra. However, current software tools often rely on single-run data for peptide peak identification, leading to inconsistent quantification across heterogeneous datasets. Match-between-runs (MBR) algorithms address this by aligning peaks or elution profiles across runs post-analysis but they are often *ad-hoc* and lack statistical frameworks for controlling peak quality, resulting in false positives and reduced quantitative reproducibility. Here we present DreamDIAlignR, a cross-run peptide-centric tool that integrates peptide elution behavior across runs with a deep learning peak identifier and signal alignment algorithm for consistent peak picking and FDR-controlled scoring. DreamDIAlignR outperformed state-of-the-art MBR methods, identifying up to 25.6% more quantitatively changing proteins on a benchmark dataset and 38.5% more on a cancer dataset. Additionally, DreamDIAlignR establishes an improved methodology for performing MBR compatible with existing DIA analysis tools, thereby enhancing the overall quality of DIA analysis.

## Introduction

The data-independent acquisition (DIA)-based liquid chromatography coupled with tandem mass spectrometry (LC-MS/MS) strategy enables accurate and reproducible protein identification and quantification for large-scale molecular biology researchPurvine et al. (2003); Venable et al. (2004). It is one of the most widely utilized high-throughput proteome profiling methods, often used for its superior performance in cross-run quantitative cohort studiesHebert et al. (2014); Aebersold and Mann (2016); Tabb et al. (2010); Bruderer et al. (2015); Ludwig et al. (2018). During the DIA data acquisition process, all peptide precursors in a relatively wide mass-to-charge (m/z) ratio window are selected for cofragmentation in an unbiased mannerChapman et al. (2014), resulting in highly multiplexed MS2 spectra that are not suitable for direct analysis by peptide search engines for the identification of peptidesKrasny and Huang (2021); Peckner et al. (2018); Bilbao et al. (2015). To address this issue, peptide-centric analysis (PECA) methodsGillet et al. (2012); Ting et al. (2015) were developed where a spectral library containing predefined m/z values, retention time (RT), fragment ion intensities and all the necessary information of the peptides of interest is used to query against raw DIA data to find the evidence of the presence of each peptide in the library at a certain confidence levelRosenberger et al. (2017). In PECA, only the chromatograms of the peptides and fragment ions in the library are extracted from the raw data and analyzed in a targeted manner, injecting strong priors into the data analysis and helping to alleviate the interference from cofragmented ions and improve identification sensitivity.

Since the concept of targeted data extraction was introduced, several software tools have been developed for PECA on DIA proteomics dataZhang et al. (2020). OpenSWATHHannes L Röst et al. (2014) pioneered automated DIA data analysis by extracting chromatograms (XICs), scoring co-eluting peak groups, and employing semi-supervised learning with FDR controlRosenberger et al. (2017); Reiter et al. (2011), establishing a foundational framework for subsequent toolsRöst et al. (2017). DIA-NNDemichev et al. (2020) built upon this foundation by incorporating additional sub-scores and leveraging a neural network model to enhance peptide identification. Taking these advancements further, DreamDIAGao et al. (2021) replaced traditional scores with a deep learning model, demonstrating superior capability in capturing complex chromatogram featuresAmodei et al. (2019).

Although existing PECA tools excel in single-run analysis, achieving consistent analyte identification and quantification across heterogeneous sample cohorts remains challengingGupta et al. (2019, 2023); Collins et al. (2017); Poulos et al. (2020). Most existing algorithms treat each run independently, without integrating information from other runsHannes L Röst et al. (2014); Demichev et al. (2020); Gao et al. (2021). When iterating over each run in a sample set one at a time, PECA tools attempt to find viable chromatographic signals, referred to as “peak groups”, for peptides in the spectral library sequentially and score these peak groups separately. However, these peak group scores reflect only the elution behavior of peptides in individual runs. Consequently, the target-decoy binary discriminative model relies solely on single-run peak group scores, overlooking the relationships among peak groups of the same peptide across all runs. This limitation can lead to inconsistent peak identification for a single peptide across different runs, especially when a high-scoring peak group originates from another peptide in one run but not in the other. Treating each LC-MS/MS run independently substantially hinders the ability of a statistical model to analyze large-scale datasets and correct for experimental idiosyncrasies in these datasetsPoulos et al. (2020). While Group-DIALi et al. (2015) attempts to leverage correlations between chromatograms across multiple runs, its effectiveness is limited to homogeneous datasets with highly similar elution signals. The fragmentation of information across runs remains a critical, unresolved challenge in computational proteomics, and overcoming it becomes essential for ensuring robust, reproducible analyses and advancing the field’s capacity to handle large-scale datasets.

To enhance cross-run peptide identification and quantification reproducibility, several match-between-runs (MBR) algorithms have been developedRöst et al. (2016); Gupta et al. (2019, 2023); Demichev et al. (2020). These MBR algorithms compare and align signals among multiple runs after regular PECA, effectively correcting the RT locations and boundaries of falsely identified peak groups *after* statistical scoring. However, despite these advances, MBR algorithms still rely on the peak groups picked and the scores assigned by single-run analysis approaches. Their performance depends on high-scoring “reference” peaks observed in one or a few runs. However, the peak with the highest single-run score across all runs may not always be the correct one due to variations in interfering signals across different runs. Consequently, if the reference peak is chosen incorrectly, the identified peaks in all runs could be completely erroneous. Moreover, MBR is typically applied after statistical scoring, and thus undermines the guarantees and safeguards provided by FDR control, potentially leading to a substantial increase of the number of false positive peak groups. Currently there are no statistical methods available capable of estimating the identification confidence of aligned peaks. In practice, MBR algorithms often require running with a heuristically increased, less strict FDR. While this approach reports more candidate peak groups, it unintentionally leads to decreased quantification performance. Consequently, MBR algorithms face a dilemma in balancing the trade-off between identification and quantification: They must either accept more low-quality but well-aligned peaks, compromising quantification accuracy, or reduce identification numbers to improve quantification.

Here, we introduce DreamDIAlignR, a cross-run peptide-centric DIA proteomics data analysis software tool. It integrates the DreamDIAGao et al. (2021) deep learning peak scorer with the MBR algorithm DIAlignRGupta et al. (2019), enabling consistent cross-run peptide identification at the entire dataset level. Instead of processing each MS injection sequentially, it considers all-run performance collectively for each library peptide. By aligning raw chromatograms first, it calculates a multi-run score for each peak, which is a weighted combination of single-run quality scores that reflects the overall performance across all runs in the dataset. Considering both single-run peak quality and multi-run global performance, the peak identification confidence can be automatically estimated using a statistical model between target peptides and decoys, eliminating the need for manual *ad-hoc* tuning of the FDR threshold. Meanwhile, it enables the discriminative model to learn the alignment relationship among peak groups across runs, thereby reducing inconsistent cross-run peak selection. In experiments conducted on both standard datasets and highly heterogeneous datasets, DreamDIAlignR substantially outperformed state-of-the-art software tools, achieving more consistent identification and enhanced quantification accuracy. Moreover, our results show that analyzing highly heterogeneous DIA data across multiple runs simultaneously yields superior cross-run protein quantification results compared to using single-run-only methods.

## Results

### Design of the DreamDIAlignR workflow

The DreamDIAlignR workflow incorporates two recent innovations in targeted DIA analysis. First, it uses a deep learning-based method to generate a continuous quality scoring profile for peptide peak groups along the RT axis in each run. Second, a dynamic programming algorithm aligns chromatographic traces, ensuring one-to-one mapping of data points across runs. These strategies seamlessly integrate peptide identification information across all runs without discontinuity or hard cutoffs. The workflow comprises the following four main steps (Figure 1b, see Methods for details).

**Figure 1.**
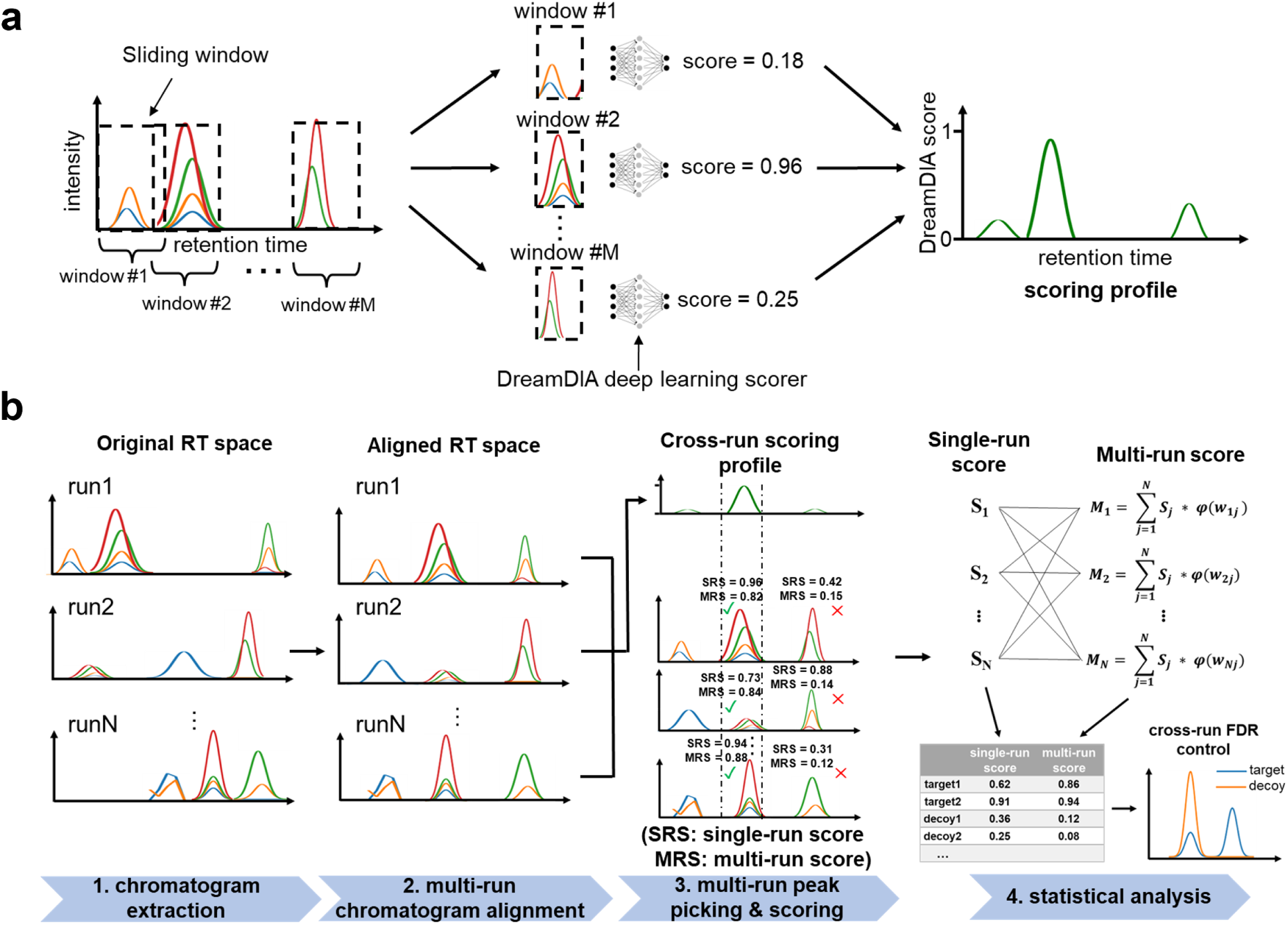
Schematic illustration of DreamDIAlignR. **a.** Working principle of the DreamDIA deep learning peak group scorer. First, a sliding window traverses the chromatogram time point by time point. Then, the signal within each window is fed into the pre-trained deep learning model to estimate its probability of being a real peptide signal (high score) or noise (low score). Finally, a continuous scoring profile is obtained for the corresponding chromatogram. **b.** DreamDIAlignR cross-run peptide-centric analysis workflow. Instead of processing individual runs sequentially, DreamDIAlignR considers all runs together for each peptide. First, DreamDIAlignR extracts chromatograms from all runs and scores them using the DreamDIA deep learning peak scorer. Next, the match-between runs (MBR) algorithm aligns the chromatograms and scoring profiles across runs. Then, the peaks are picked based on a majority voting of multiple runs represented by an averaged cross-run scoring profile instead of relying on a reference peak, and the multi-run scores are calculated by aggregating scores of corresponding peaks across runs. Finally, DreamDIAlignR considers both single-run scores and multi-run scores for statistical analysis.

#### Chromatogram extraction and scoring

DreamDIAlignR first extracts chromatograms for precursor and fragment ions across all runs for each library peptide. It then uses a pre-trained Long Short-Term Memory (LSTM) modelHochreiter and Schmidhuber (1997); Gao et al. (2021) to slide a scoring window along the RT, generating continuous scoring profiles that estimate the probability of each time point and its context representing a true peptide peak group (Figure 1a).

#### Multi-run chromatogram alignment

To address RT misalignment caused by sample heterogeneity and experimental variationsNigjeh et al. (2017), we then employ MBR algorithms to synchronize chromatograms and scoring profiles across runs, including both run-wide global alignmentSmith et al. (2015); Röst et al. (2016) and peptide-wide dynamic alignment via DIAlignRGupta et al. (2019, 2023) to account for peptide-specific elution behavior.

#### Multi-run peak picking and peak scoring

By averaging aligned single-run scoring profiles, DreamDIAlignR generates a cross-run scoring profile that represents the collective elution behavior of each peptide across all runs. Candidate peak groups with the highest averaged scores are then picked, ensuring identification is based on majority consensus rather than relying solely on single-run dataHannes L Röst et al. (2014). Simultaneously, we introduce a multi-run score for each peak group, which combines single-run scores weighted by global RT similarity, providing a balanced assessment of both individual and cross-run elution behavior to improve peak group identification.

#### Statistical analysis

The output of DreamDIAlignR is a discriminative model to distinguish between real peptides and artificially created decoysReiter et al. (2011); Rosenberger et al. (2017); Demichev et al. (2020). In contrast to routine methods that only concern peak groups’ single-run behavior, DreamDIAlignR extends existing statistical approachesReiter et al. (2011); Rosenberger et al. (2017) to learn the peak group correspondence across runs by utilizing both single-run scores and multi-run scores, thereby enabling more consistent peak selection and more comprehensive confidence estimation with a cross-run horizon.

### Feasibility of cross-run signal alignment and integration

We first performed a feasibility test of our cross-run signal alignment and integration strategy using a pilot dataset, the *S. pyogenes* datasetRöst et al. (2016); Gupta and Röst (2021), which included approximately 7000 manually annotated peak groups across 16 LC-MS/MS injections (Supplementary Figure S10a).

As an example, the DreamDIA scoring profile for each run of the target peptide (Figure 2b) is obtained by moving the DreamDIA peak group scorer along the RT axis of the corresponding chromatograms (Figure 2a). Notably, the score apex regions show major concordance with the manually annotated peak group regions, indicating the peak identification accuracy of the DreamDIA scorer. In addition, although RT shifts among peak groups across multiple runs are clearly visible, the signals can be effectively synchronized when MBR algorithms like DIAlignR are applied (Figure 2c). While all the MBR algorithms mitigate RT discrepancies across multiple runs, the chromatograms aligned by either global lowess method or DIAlignR demonstrate superior synchronization compared to the global linear method (Supplementary Figure S1).

**Figure 2.**
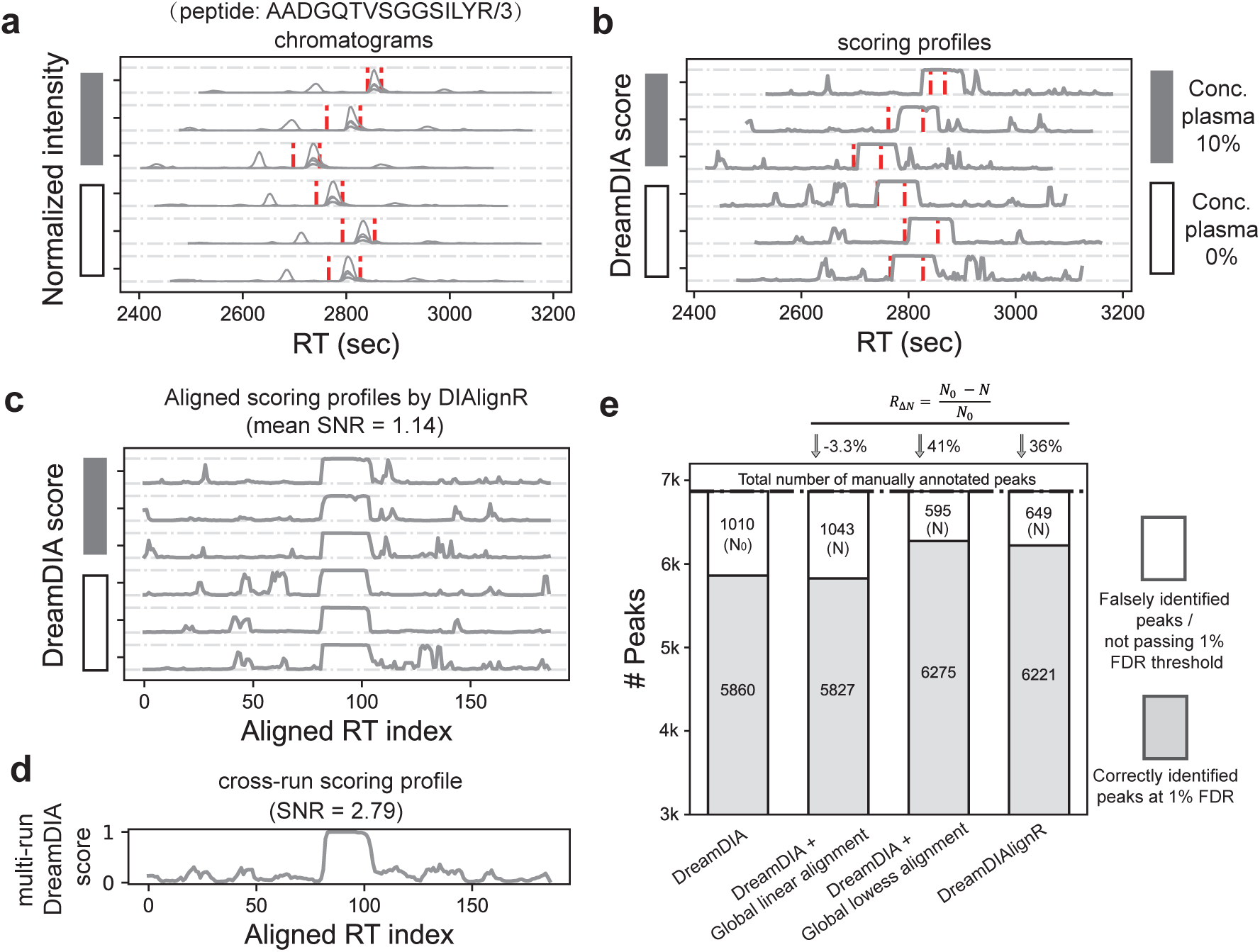
Chromatogram alignment and integration methods in the DreamDIAlignR workflow and their performance on a *Streptococcus pyogenes* (*S. pyogenes*) dataset. a-d. Intermediate results of an example peptide “ADGQTVSGGSILYR/3” in the *S. pyogenes* dataset being processed by DreamDIAlignR. Results of 6 out of 16 runs are shown due to limited space. **a.** Chromatograms of 500-1000 seconds wide are extracted in all runs. Red dashed lines denote manually annotated peak boundaries. **b.** A continuous scoring profile is calculated for each run by DreamDIA deep learning peak group scorer. **c.** Aligned scoring profiles of all runs by DIAlignR. **d.** Averaged scoring profile of all runs. Signal-to-noise ratio (SNR) is calculated as the maximum score of manually annotated peak regions divided by the maximum score of the other regions. **e.** Comparison of the number of correctly and incorrectly identified peak groups using different signal alignment methods. The incorrect identification number of DreamDIA without any alignment strategy is regarded as a baseline (*N*_0_), which is used to calculate the decreasing rates (*R*_Δ_*_N_*) when applying signal alignment algorithms.

Moreover, although DreamDIA can correctly identify the locations of peak groups, there still remains some high-score noisy signals in the scoring profiles that fall outside the true peak group regions (Figure 2b, c). To address this, we averaged the aligned scoring profiles from multiple runs. This process diluted and suppressed the random noise originating from a single experiment alone, resulting in a cross-run scoring profile with over 2-fold higher signal-to-noise ratio (SNR) compared to the single-run profiles (Figure 2d). The averaged scoring profile will then be utilized for cross-run peak picking, which yields greater accuracy and consistency compared to sequential single-run peak picking methods.

We next evaluated the accuracy of peak group identification using manual annotations as ground truth, comparing different MBR approaches and DreamDIA without multi-run analysis (Figure 2e). Compared to regular DreamDIA, global lowess alignment and DIAlignR reduced false identifications by 41% and 36%, respectively, while global linear alignment showed no improvement. This aligns with previous findingsGupta et al. (2019), where global linear models failed to address run-to-run RT variation for this dataset. Non-linear lowess alignment also achieved lower residuals than linear alignment, demonstrating its robustness for datasets with significant RT shifts (Supplementary Figure S2). DIAlignR, on the other hand, directly aligns chromatograms across runs for individual peptides. It provides the flexibility to adjust the alignment function for each peptide, resulting in comparable results to the global lowess alignment. It is noteworthy that different datasets exhibit varying cross-run retention time discrepancy patterns indicating that the performance of various MBR algorithms can vary across datasets. When choosing and benchmarking MBR algorithms, flexibility and robustness play a crucial role.

### Simultaneously improved identification and quantification performance

We benchmarked DreamDIAlignR’s peptide and protein identification and quantification performance using the LFQbench HYE110 dataset, which includes replicated two samples with known inter-species abundance ratios (human 1:1, yeast 10:1, and *E. coli* 1:10) for evaluating DIA tools’ ability to recover these ratiosNavarro et al. (2016) (Supplementary Figure S10b).

We first evaluated the MBR algorithms of DIA-NN, OpenSwath and DreamDIAlignR at 1% FDR (Figure 3a and Supplementary Figure S3a). DreamDIAlignR uniquely improved both identification and quantification, identifying 27.0% more peptides and reducing quantification bias by 12.7%. In contrast, DIA-NN’s MBR increased peptide identification by 11.9% but raised quantification bias by 53.5%. OpenSWATH’s MBR reduced quantification bias by 27.8% but at the expense of a slight 3.0% loss in peptide identification. Benchmarking at the protein level showed similar trends (Supplementary Figure S3a). These findings indicate that both OpenSWATH and DIA-NN have to trade-off quantitative accuracy with identification performance during MBR (with DIA-NN producing more, but quantitatively worse peptide identifications while OpenSWATH produces less, but quantitatively better peptide identifications). However, this is not the case for DreamDIAlignR which achieved better results in both, identifying 21.5% more peptides than OpenSWATH and 10.2% more than DIA-NN, with quantification bias 47.7% and 27.8% lower, respectively.

**Figure 3.**
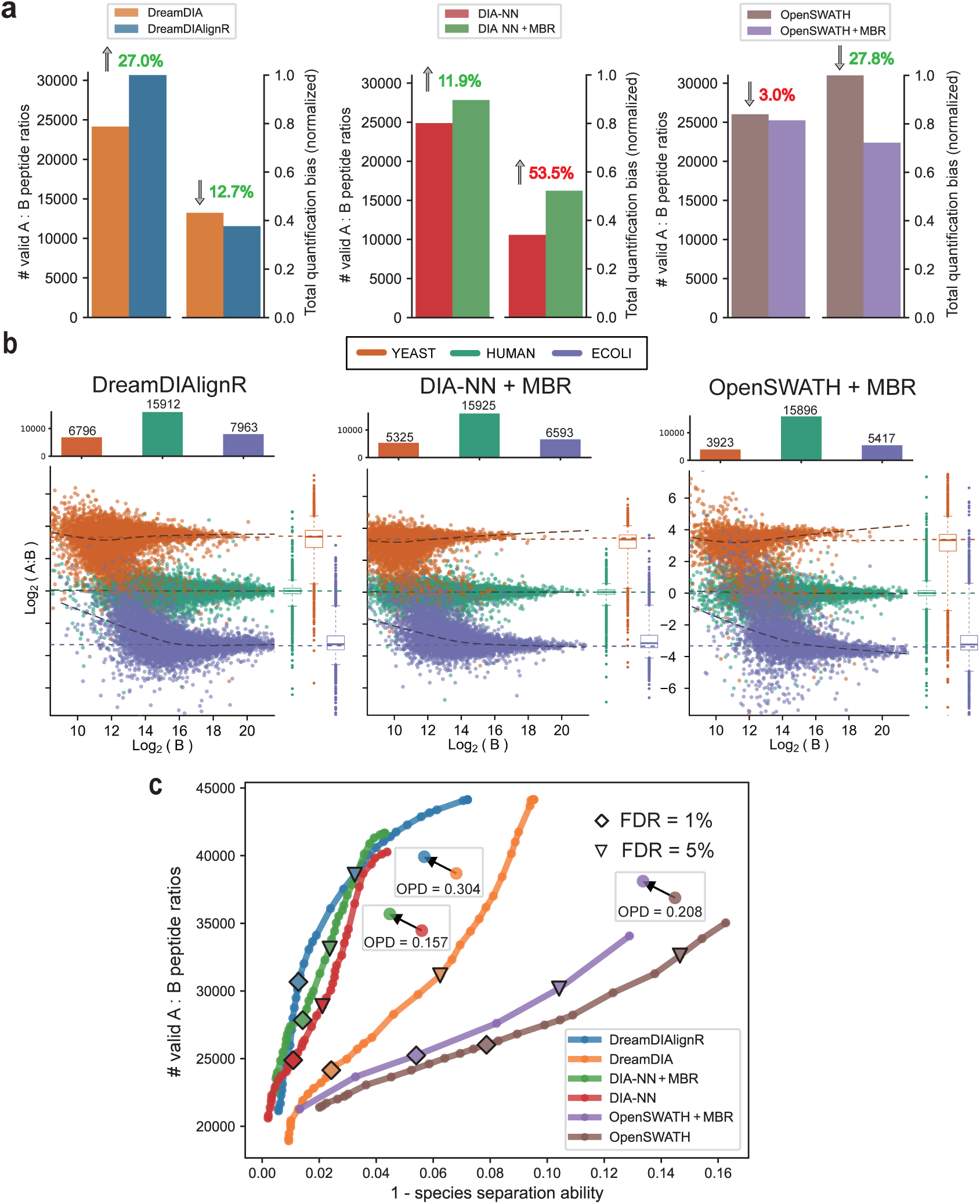
Identification and quantification performance benchmark on LFQbench dataset. **a.** The number of valid peptide precursor ratios identified and the total quantification bias of various software tools before and after applying match-between runs at 1% precursor FDR. Total quantification bias is calculated as the mean of the three normalized quantification performance metrics given by LFQbench software suite including 1 - species separation ability, median bias and dispersion (See Methods). **b.** Peptide-level LFQbench benchmark results of OpenSWATH + Match-Between-Runs (MBR), DIA-NN + MBR and DreamDIAlignR at 1% precursor FDR. Colored dashed lines denote log-transformed ground truth Sample A to Sample B ratios (Human, 1:1; Yeast, 10:1; *E. coli*, 1:10). The numbers of valid peptide ratios identified for different species are shown in the bar charts. Boxplot elements: center line, median; boxes, interquartile range; whiskers, 1.5x interquartile range; points, outliers. **c.** The number of valid Sample A to Sample B peptide precursor ratios identified in total and the corresponding 1 - species separation ability for quantification using a series of FDR thresholds. The species separation ability, as provided by LFQbench software package, represents the area under the receiver operating characteristic (ROC) curve of a binary classifier between two species. Here we show the mean of the 1 - species separation ability of human, yeast and *E. coli* peptides. “♢” and “▽” mean the results at 1% and 5% peptide precursor FDR respectively. The Overall Performance Distance (OPD) evaluates the improvement in identification and quantification performance of each software tool, comparing results before and after the application of MBR (See Methods).

In many experimental settings, accurate identification and quantification of analytes that change in abundance are of particular interest since these could indicate proteins of interest for a drug target, biomarker or mechanistic study. We therefore focused our analysis next on the yeast and *E. coli* proteomes which comprise the variable part of the A:B mixture experiment. DreamDIAlignR showed superior cross-run quantification accuracy compared to OpenSWATH and DIA-NN, with the A:B quantification ratios closer to the ground truth ratio line for each species and slightly higher dispersion than DIA-NN (Figure 3b and Supplementary Figure S3b). Notably, all software tools yielded comparable numbers of human peptides and proteins, while DreamDIAlignR outperformed the other tools in identifying yeast and *E. coli* analytes. It improved yeast and *E. coli* peptide precursor identifications by 27.6% and 20.8%, respectively, over DIA-NN, and by 73.2% and 47.0%, respectively, over OpenSWATH. Protein-level improvements were 25.6% and 18.7% over DIA-NN and 58.6% and 39.3% over OpenSWATH. Manual inspection confirmed superior cross-run peak group identification consistency and quantification accuracy, particularly in low-abundance runs (Supplementary Figure S6). These findings highlight DreamDIAlignR’s ability to detect up to 58% more quantitatively changing proteins without compromising accuracy, offering significant advantages for multi-run analyses.

To negate biases in over-optimistic FDR control by one tool, we next analyzed our results across a large range of FDR cutoffs, allowing us to evaluate the overall performance more accurately by minimizing the influence of fixed FDR thresholds. To intuitively demonstrate the improvement introduced by MBR methods, we also introduced a metric named Overall Performance Distance (OPD) (Figure 3c). OPD quantifies the “distance” between the overall performance of a software tool before and after applying MBR. This measurement is based on a combination of four factors: the number of valid peptide ratios, 1 - species separation ability, median bias, and dispersion. The first factor pertains to identification, while the remaining three address quantification. These four metrics collectively form a 4-dimensional point representing the software tool’s overall performance at a specific FDR threshold. The OPD is then calculated as the median Euclidean distance between each pair of metric points at the same FDR threshold, before and after the application of MBR (See Methods). The results indicated that, without considering hard FDR cutoffs, OpenSWATH, DIA-NN, and DreamDIA all exhibited effective overall performance improvements after applying MBR (Figure 3c and Supplementary Figure S4). Specifically, the OPDs revealed that DreamDIAlignR’s MBR method (OPD = 0.304) achieved a 94.4% greater overall performance improvement compared to DIA-NN (OPD = 0.157), and a 46.4% improvement compared to OpenSWATH (OPD = 0.207). These results showed that DreamDIAlignR substantially improved MBR performance and achieved the best overall performance among all tools, with or without the MBR strategy. Additionally, benchmarking various MBR algorithms available in DreamDIAlignR revealed that the DIAlignR algorithm outperformed the other alignment methods (Supplementary Figure S5).

### The multi-run score ensures reliable cross-run peptide identification

Statistical error control is essential for ensuring reliable multi-run proteomics analysisRosenberger et al. (2017). Lim et al.Lim et al. (2019) highlighted this, showing that MBR algorithms for DDA data caused an 8-fold increase in false identifications, including the erroneous identification of yeast peptides in human-only samples in their 2-Sample, 2-Proteome ChallengeYu et al. (2021). To mitigate erroneous alignment, DreamDIAlignR employs an exponential weight decay function governed by a penalty parameter *k*, which calculates multi-run scores by assigning higher weights to similar runs while penalizing contributions from distant ones. This approach prioritizes relevant scoring information, minimizing false identifications and avoiding inflated scores, particularly in highly heterogeneous datasets.

Herein, we investigated the impact of the parameter *k* on the identification performance. A subset of 24 samples, consisting of 12 human-proteome-only runs and 12 human-yeast mixed runs from the Procan large-scale cancer studyPoulos et al. (2020), were selected for testing (Supplementary Figure S10c). As *k* increased, the number of peptides identified declined, with a watershed *k* value of 50 (Figure 4a). Compared to DreamDIAlignR without weight decay (*k* = 0), the number of yeast peptides falsely identified in the human-only runs decreased by 92.7% when *k* = 50. This reduction was significantly greater than the decreases observed in the yeast peptides identified in human-yeast mixed runs (11.7%) and human peptides in human-only runs (4.2%). We also monitored changes in FDR before and after applying the weight decay function. Two-species FDRs were calculated both with and without accounting for the differing likelihoods of human and yeast peptide identification, represented as the upper-bound and lower-bound FDRsWen et al. (2024), respectively (See Methods). With a *k* value of 50, the upper-bound two-species FDR significantly decreased from 16.4% to a much more acceptable 1.28%, while the lower-bound FDR also dropped from 3.46% to 0.27%. These findings indicate that the weight decay function effectively alleviates false positive identifications while only minimally impacting true positive ones, thereby ensuring reliable cross-run FDR control.

**Figure 4.**
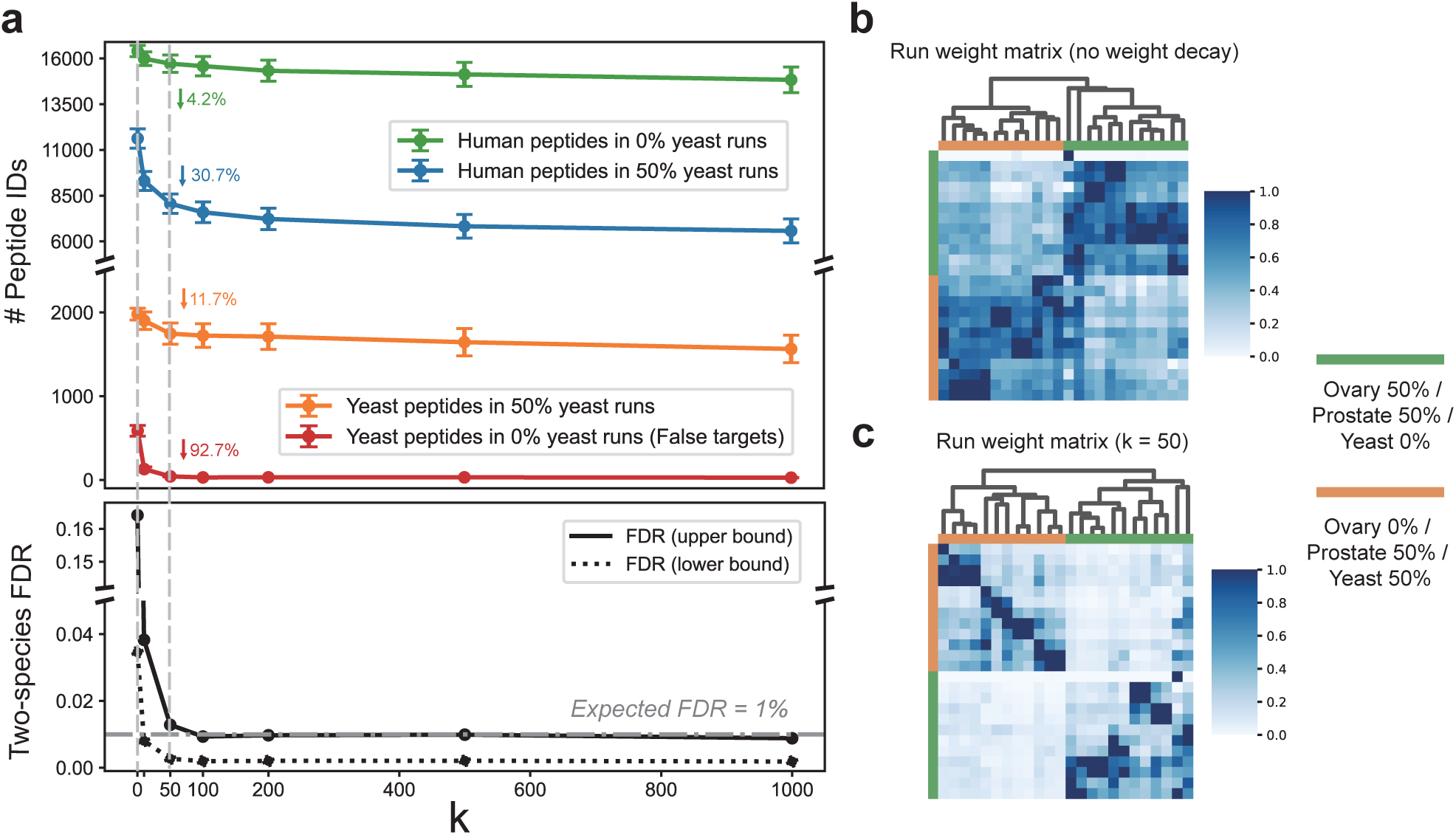
Optimization of weight decay parameters for multi-run score calculation on the Two-Sample, Two-Proteome (TSTP) dataset. **a.** The number of peptide precursors identified and the corresponding two-species FDR, calculated from mixed and human-only samples using various weight decay parameters, denoted as *k*. The parameter *k* controls the penalization of distant runs in the exponential weight decay function. A *k* value of 0 indicates that the weight decay approach has been deactivated. The number of yeast peptides identified in human-only runs serves as a measure of false targets. The two-species FDR was calculated using the “combined” method reported by Wen et alWen et al. (2024) (See Methods). **b.** Run weight matrix without weight decay, showing hierarchical clustering based on similarity metrics. **c.** Run weight matrix with weight decay (*k* = 50), showing hierarchical clustering based on similarity metrics.

In DreamDIAlignR, the weights of different runs in the multi-run score of a target run are determined by their global RT similarity to the target run. This ensures that a low-quality peak will not receive an undesirably high multi-run score, even if it has been aligned to high-quality peaks in other runs, due to the low similarity between the runs. To confirm that the scoring information from different runs is correctly weighted, we inspected the run weight matrices before (Figure 4b) and after (Figure 4c) applying weight decay. Our observations indicate that, in this dataset, sample replicates (considered highly similar runs) contributed more significantly to the multi-run scores, while distant runs (runs of different samples) were largely ignored. In addition, the multi-run score, which indicates the overall quality of the aligned peaks across all runs, is a novel concept that current PECA software tools do not possess. Therefore, we investigated whether the multi-run score could enhance the discriminative power of the statistical model to distinguish between target peptides and decoys. Results showed that the multi-run scores of yeast peptides in human-yeast mixed runs (true targets) were significantly higher than those in human-only runs (false targets) after applying weight decay (Supplementary Figure S7a, b). Moreover, the multi-run score ranked second among all the sub-scores used by the statistical model, with a feature importance of 27.0% (Supplementary Figure S7c). An ablation test also showed that discarding all multi-run scores before building the statistical model caused the number of identifications to revert to levels comparable to DreamDIA (Supplementary Figure S7d). These results highlight that the multi-run score significantly aids the discriminative model in accurately identifying yeast peptides in the appropriate runs.

Note that the dataset used in this experiment simulates an extreme scenario of biologically heterogeneous data, where the two samples have completely different compositions. In actual biomedical studies, it is uncommon to align human runs with yeast runs. Therefore, the penalty parameter optimized in this experiment might be overly conservative. To benchmark our tool from the most conservative perspective possible, we applied *k* values of 50 or higher in all the experiments in this work. Researchers are able to select a *k* value less than 50 in their own proteomics studies to achieve improved performance.

### Better identification and quantification performance for highly heterogeneous datasets

One notable feature of DreamDIAlignR is its ability to analyze highly heterogeneous datasets through signal alignment and integration of multiple runs. Therefore, we evaluated its performance on the ProcanPoulos et al. (2020) cancer dataset, featuring mixed-species proteomes similar to LFQbench (Ovary 1:4, Prostate 1:1, Yeast 1.75:1) but with greater heterogeneity due to acquisition on different instruments over extended intervals (See Methods, Supplementary Figure S10d).

At a 1% precursor FDR, DreamDIAlignR remained the only MBR approach that simultaneously improved both identification and quantification performance. Compared to DreamDIA single-run analysis, DreamDIAlignR increased the number of identified peptides by 168.6% while reducing quantification bias by 1.6% (Figure 5a). In contrast, OpenSWATH maintained a similar number of identifications but primarily reduced quantification bias by 21.6% after applying MBR. DIA-NN increased identification numbers by 24.5%, but this came at the expense of increasing quantification bias by 25.8%. Among all the software tools utilizing the MBR strategy, DreamDIAlignR achieved the highest number of identifications and the highest quantification accuracy simultaneously. We also calculated the OPDs to assess the overall performance improvement brought by different MBR methods, independent of FDR threshold selection. The overall identification and quantification improvement of DreamDIAlignR (OPD = 0.368) was 292.9% greater than that of DIA-NN (OPD = 0.094) and 764.7% greater than that of OpenSWATH (OPD = 0.043) (Figure 5b, c). These results demonstrate that DreamDIAlignR significantly outperforms current data analysis tools and MBR approaches, particularly for highly heterogeneous data. In addition, among the three MBR algorithms available in DreamDIAlignR, DIAlignR remained the best for this dataset with lower median bias compared to global alignment methods (Supplementary Figure S8). With significantly more analytes identified, we then investigated whether DreamDIAlignR provided a more complete quantification matrix. The results showed that DreamDIAlignR improved data completeness by 12.0% at the peptide precursor level (Figure 5d) and 7.7% at the protein level (Figure 5e) compared to DreamDIA single-run analysis. Furthermore, among all software tools, both with and without MBR strategies, DreamDIAlignR achieved the highest data completeness—4.0% and 6.2% higher than DIA-NN with MBR and OpenSWATH with MBR, respectively, at the peptide precursor level, and 3.8% and 5.1% higher at the protein level.

**Figure 5.**
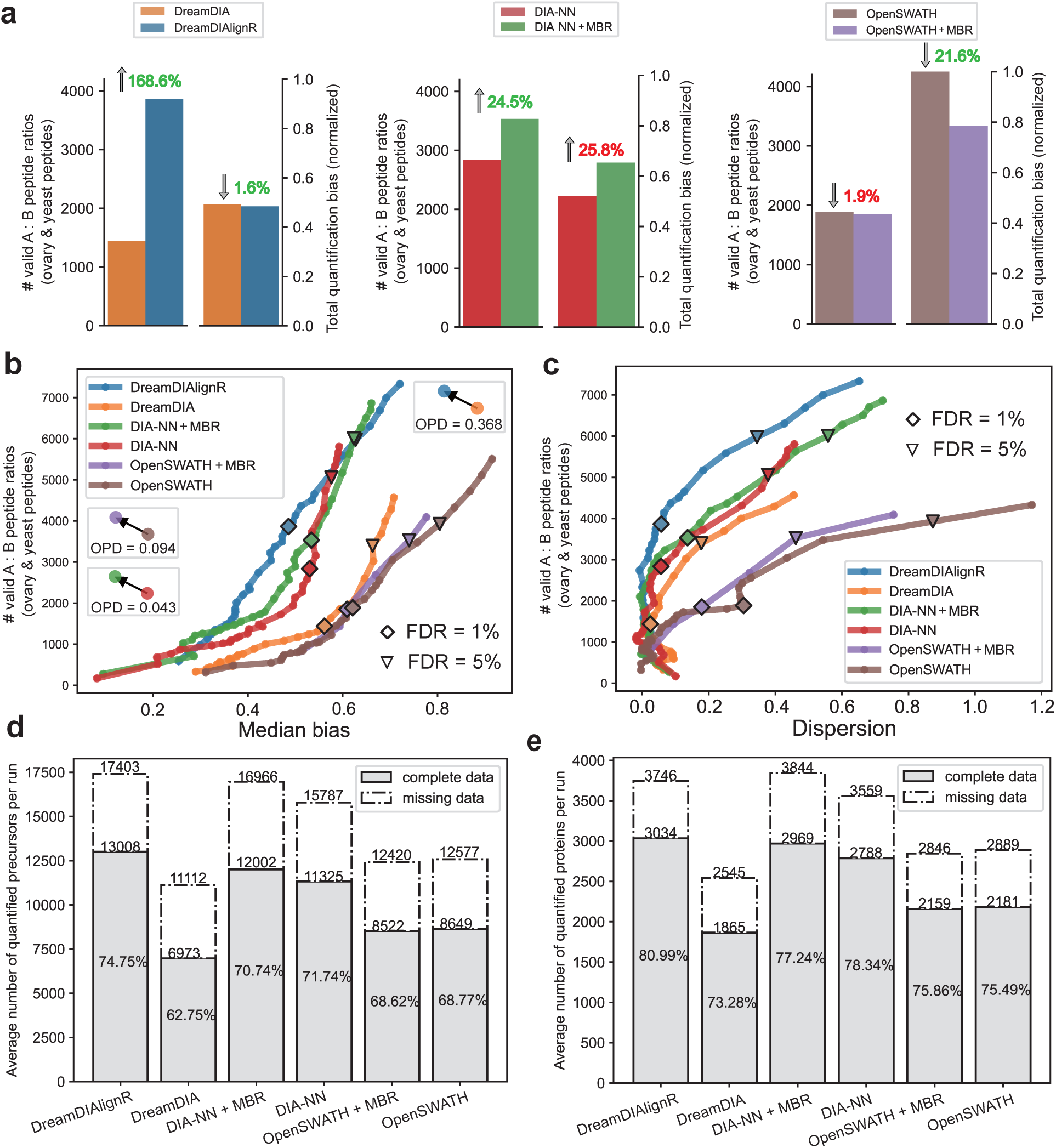
Identification and quantification performance benchmark on highly heterogeneous dataset. **a** The number of valid peptide precursor ratios identified and the total quantification bias of various software tools before and after applying match-between runs at 1% precursor FDR. Total quantification bias is calculated as the mean of the two normalized quantification performance metrics: median bias and dispersion. **b, c.** The number of valid yeast and ovary Sample A to Sample B peptide precursor ratios is plotted against the corresponding median bias (**b**) and dispersion (**c**) using a series of FDR thresholds. A ratio is considered valid only if a peptide is identified in at least 3 Sample A runs and 3 Sample B runs. Quantification bias metrics are calculated as recommended by the LFQbench software package. The median bias is defined as the distance between the median of quantified Sample A to Sample B ratios and the ground truth ratio lines. Here we show the mean of the median biases for ovary-specific and yeast peptides. Dispersion is defined as the standard deviation of the log-transformed ratios for each species. We also present the mean dispersion for ovary-specific and yeast peptides. “♢” and “▽” mean the results at 1% and 5% peptide precursor FDR respectively. The Overall Performance Distance (OPD) evaluates the improvement in identification and quantification performance of each software tool, comparing results before and after the application of MBR (See Methods). **d, e.** Benchmark of quantification matrix completeness at the peptide precursor level (**d**) and the protein level (**e**). The percentages indicate the ratios of validly quantified peptide precursors or proteins to the total number of analytes identified. Solid-lined bars represent the average number of quantified peptide precursors or proteins across all runs, while dashed-lined bars represent the total dimension of the quantification matrix divided by the total number of runs (36). The quantification matrices were filtered with a 1% precursor FDR, and peptides identified in fewer than 3 out of 36 runs were excluded.

Moreover, to evaluate DreamDIAlignR on large-scale datasets, we analyzed the Procan494 dataset, consisting of 494 runs with technical replicates and varying species concentration ratios, enabling calibration curve generation (Supplementary Figure S10e). Compared to DIA-NN, DreamDIAlignR demonstrated superior identification and quantification, identifying 21.2% more peptides and achieving a 2.7% higher median R² at 1% FDR (Supplementary Figure S9). This result highlights the robust performance of DreamDIAlignR in large-scale, high-throughput proteome profiling.

### Improved cross-run quantification provides more insightful data for biological analysis

Drawing meaningful biological conclusions in cohort proteomics studies hinges on the precise identification of differentially expressed signature proteins across samples. Achieving this requires robust and accurate cross-run peptide quantification, which enables the reliable detection of significant fold changes through statistical analysis amidst a large background of non-changing signals. Given that DreamDIAlignR has demonstrated superior cross-run quantification performance in benchmarking studies, we seek to evaluate its potential for providing deeper insights into biological changes in proteomic analyses.

We conducted differential expression analysis on the 2-Sample, 2-Proteome dataset using limmaRitchie et al. (2015). Since only the Sample B runs contain ovarian cancer cells (Supplementary Figure S10c), the up-regulated proteins in these runs can be considered ovarian cancer-related proteins. Our results demonstrated that DreamDIAlignR identified 38.5% and 112.4% more differentially expressed proteins at p < 0.01 using limma compared to DIA-NN with MBR and OpenSWATH with MBR, respectively (Figure 6a). While a substantial overlap was observed in the proteins identified by the different software tools, DreamDIAlignR detected a greater number of unique proteins that are quantitatively changing (Figure 6b). This suggests that DreamDIAlignR has a higher potential to uncover meaningful functional proteins or biomarkers in cohort-based proteomics studies.

**Figure 6.**
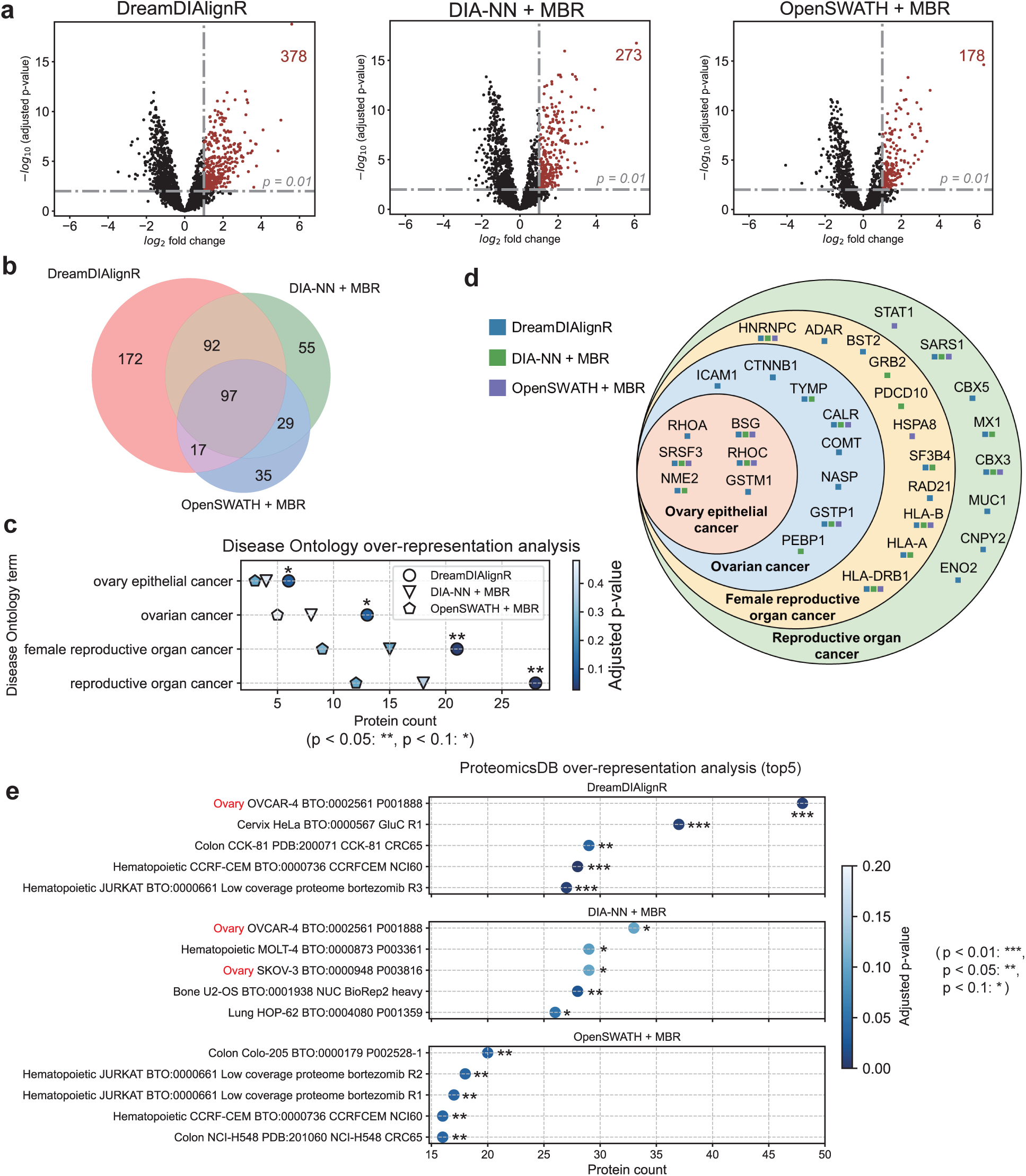
Comparison of differential expression analysis on the Two-Sample, Two-Proteome (TSTP) dataset conducted by three match-between-runs (MBR)-enabled software tools. **a.** The numbers of differentially expressed proteins identified by DreamDIAlignR, DIA-NN with MBR and OpenSWATH with MBR in the 50% ovarian cancer tissue runs. The horizontal dashed lines indicate an adjusted p-value threshold of 0.01. The vertical lines represent a fold-change cut-off of 2. Proteins outside these thresholds are considered significantly up-regulated in the runs with 50% ovarian cancer tissue. **b.** Consistency comparison of differentially expressed proteins identified by various software tools. **c.** Over-representation analysis using Disease Ontology (DO) for the up-regulated proteins shown in **a**, highlighting DO terms related to ovarian cancer. **d.** Ovarian cancer-related proteins identified in the DO analysis using three different software tools. **e.** Comparison of the top 5 over-represented gene sets in ProteomicsDB for the up-regulated proteins, identified using different software tools.

To assess the biological relevance of the increased protein identifications by DreamDIAlignR, we performed over-representation analysisXu et al. (2024) on the up-regulated proteins from the Sample B runs using two databases: Disease Ontology (DO)Schriml et al. (2022) and ProteomicsDBSamaras et al. (2020). In the DO analysis, the up-regulated proteins identified by DreamDIAlignR showed a stronger over-representation in ovarian cancer-related DO terms and yielded significantly lower p-values compared to the results from the other two software tools (Figure 6c). Consistent with the overall overlap in differentially expressed proteins (Figure 6b), DreamDIAlignR detected the majority of ovarian cancer-related proteins identified by the other two software tools, with its uniquely identified proteins also showing a strong association with ovarian cancer (Figure 6d). Manual inspection of peaks identified by various software tools further showed that DreamDIAlignR delivered more complete and accurate cross-run quantification while effectively filtering out low-quality peaks (Supplementary Figures S11 and S12). This capability enhances differential expression analysis, providing higher statistical confidence compared to other tools. Since Disease Ontology serves as a general semantic database primarily focused on linking genomic data with disease features and mechanismsSchriml et al. (2022), we additionally utilized ProteomicsDBSamaras et al. (2020)—a database providing orthogonal disease information represented at the protein level—to perform over-representation analysis from a proteomics perspective. To ensure the completeness of the analysis results for different software tools, here we set the p-value cutoff for the over-representation analysis to 0.1 (while maintaining a 1% precursor FDR cutoff for all peptide identifications). Our results indicate that both DreamDIAlignR and DIA-NN with MBR rank OVCAR-4, a human ovarian cancer cell line, as the top gene set in the over-representation analysis (Figure 6e). Notably, DreamDIAlignR identifies 45.5% more OVCAR-4 proteins than DIA-NN and achieves a significantly lower p-value (below 0.01). If the p-value cutoff is set to 0.5 or stricter, DreamDIAlignR remains the only tool capable of ranking the ovarian cancer cell line gene set first. Our analysis reveals that rather than merely improving the number of identifications, DreamDIAlignR strengthens DIA data analysis by enhancing the accuracy and coverage of identifying quantitatively changing analytes across samples. This capability supports the discovery of biologically meaningful signature proteins, providing valuable insights in real biomedical studies that involve highly heterogeneous data.

## Discussion

Consistently identifying and accurately quantifying peptides in biologically or technically heterogeneous samples remains not only the ultimate goal in DIA-based high-throughput proteomics research, but also a highly demanding task for all DIA data analysis toolsGao et al. (2021). Among the challenges encountered by DIA data analysis tools, a prominent one stems from the increased stochasticity of peptide elution signal selection across multiple samples, particularly when these samples have highly varied biological compositions or data acquisition conditions. Hence, the concept of harnessing multiple-run signal to alleviate interference and collectively identify chromatographic peaks is intuitive and has also been exploredLi et al. (2015); Yan et al. (2023); Zhang et al. (2016). However, these existing tools were developed with the assumption of data homogeneity and only incorporated rudimentary signal alignment approaches, which were not suited for highly heterogeneous data. As a result, their use has been significantly limited in large-scale biomedical studies. To address this, current MBR algorithmsRöst et al. (2016); Gupta et al. (2019, 2023) prioritized fixing the retention time shift in heterogeneous data after a single-run analysis routine. However, they leave users with a dilemma: whether to accept aligned peaks of potentially lower quality or whether to reject potentially correct peaks to increase the stringency, due to the absence of a statistical framework for MBR. In essence, current “multi-run analysis tools” fall short of true multi-run capabilities, still confined by the conventional single-run analysis perspective.

In this study, we present a novel approach for performing MBR in DIA, which integrates data from all available LC-MS/MS runs and applies statistical error control *after* the match-between-runs. This novel concept is in principle applicable to all DIA analysis tools such as DIA-NN, OpenSWATH or SpectronautBruderer et al. (2015), and addresses one of the last remaining challenges of DIA analysis: handling large-scale, heterogeneous study designs. Our findings demonstrate that DreamDIAlignR stands out as the only MBR method capable of simultaneously improving both identification and quantification performance, achieving a 38.5% increase in differentially expressed proteins in a realistic case-control dataset. As LC-MS/MS technology and throughput continue to improve, we anticipate that DreamDIAlignR will not only assist proteomics researchers but also provide deeper insights into the analysis of large-scale heterogeneous omics data.

## Methods

### DreamDIAlignR workflow

DreamDIAlignR was developed based on DreamDIAGao et al. (2021) and DIAlignRGupta et al. (2019) software libraries. The workflow of DreamDIAlignR includes four major steps described (See Figure1b) as follows. ***chromatogram extraction and scoring.*** Before extracting chromatograms, DreamDIAlignR first performs RT normalization using endogenous peptides sub-sampled from spectral libraries. For each run, DreamDIAlignR identifies the RT locations with the highest scores for all the endogenous iRT peptides across the entire RT gradient using a pre-trained deep learning peak group scorer. This scorer is the built-in LSTM deep learning model in DreamDIAGao et al. (2021), used without re-training or parameter fine-tuning. The RT location with the highest score is designated as the best RT. Peak groups located at these best RTs, with spectral cosine similarity scores above a certain threshold (0.95 by default), are considered validly identified iRT peptides. These best RTs are then used to fit a linear or non-linear model against their corresponding iRT values in the library. In DreamDIAlignR, the RT normalization step serves to both fit an RT versus iRT model to narrow down the peak group searching rangeEscher et al. (2012); Bruderer et al. (2016); Hannes L Röst et al. (2014) and to obtain the global similarity among all runs. iRT peptides with fitting residuals below a specified threshold in each run are deemed confidently identified, or inliers. The NC similarityGupta et al. (2023) of these inlier sets between each run pair in the dataset is calculated as:

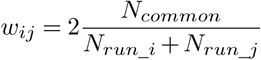

where *w_ij_* denotes the NC similarity, which is regarded as the global RT similarity between run pairs, and *N_common_* represents the number of common inlier peptide IDs in both *run_i_* and *run_j_*. The NC similarities of all run pairs collectively form a global similarity matrix for all runs.

Subsequently, DreamDIAlignR extracts the chromatograms for the top 6 fragment ions by default for all the target peptides and decoys in each run. With extracted chromatograms as input, the deep learning scorer slides along the RT ranging 200-1500 seconds, and scores the signals in each sliding window as a candidate peak group. To enhance the robustness and transferability of our approach, the scoring model also incorporates theoretical fragment ions based on the amino acid sequences of the peptides for peak group identificationGao et al. (2021). Consequently, both archived fragment ions from libraries and theoretical fragment ions have their respective chromatograms extracted in this process. DreamDIAlignR then calculates three additional single-run scores for each peak group except the DreamDIA deep learning score: spectral cosine similarity score, MS1 area score and MS2 area score. All these four scores create continuous traces along the RT axis, named as scoring profiles. The XICs and scoring profiles for each run are stored in SQLite database files for future use.

#### Multi-run chromatogram alignment

Due to sample heterogeneity and variations in experimental conditions across acquisitions, the XICs and scoring profiles of different runs often exhibit RT misalignmentNigjeh et al. (2017). To make the chromatograms and scoring profiles of multiple runs comparable, we introduced MBR algorithms including global RT alignment (run-wide) and DIAlignR alignment (peptide-wide). The global alignment strategy assumes a systematic RT variation between run pairs, where the elution order of peptides is preservedSmith et al. (2015); Röst et al. (2016). In DreamDIAlignR, a subset of peptides from the spectral library is initially selected as anchor peptides to build RT alignment models. For each pair of runs, a global RT model is constructed using the best RTs of commonly and confidently identified anchor peptides. DreamDIAlignR provides both linear and lowess modeling options to capture RT correspondence between run pairs, enabling alignment of chromatograms for each peptide across all runs. However, this approach might smooth out peptide-specific RT shifts and lead to false identifications if the elution order of peptides differs across runsSmith et al. (2015); Spicer et al. (2010); Wu et al. (2016). Therefore, we further introduced peptide-wide DIAlignR alignment that can be optionally applied on top of global alignment constraint. DIAlignR aligns chromatograms for each peptide between two runs, but its time complexity, *O*(*N_peptides_* × *N^2^_runs_*), can become prohibitive if all run pairs are aligned, especially in large sample cohorts. To address this, we constructed a minimum spanning tree based on the global similarity matrix, ensuring alignment only between run pairs connected by the tree’s edgesGupta et al. (2023). This approach reduces the time complexity to *O*(*N_peptides_* × *N_runs_*) and confines signal alignment to directly connected, highly similar runs, thereby minimizing false alignments and preventing information transfer between highly heterogeneous runs. After aligning chromatograms across multiple runs, the aligned RT vectors are interpolated and collapsed to produce a synchronized RT matrix for all runs. Users can flexibly choose between global RT alignment alone or the hybrid DIAlignR alignment. Benchmark results indicate that while the global RT alignment is more time-efficient, DIAlignR achieves higher accuracy.

#### Multi-run peak picking and peak scoring

With the RT of all runs aligned, the single-run scoring profiles are also synchronized. We then average the single-run scoring profiles across all runs to get the cross-run scoring profile. The top 10 candidate peak groups across all runs with the highest averaged deep learning scores are picked. This approach guarantees uniform peak selection and enhances comparability across the dataset.

The multi-run scores are then calculated for each candidate peak group. For each run, the multi-run scores are calculated as a linear combination of the corresponding single-run scores in all runs (Figure 1b). As a result, we can also obtain four multi-run scores for each peak group as counterparts for the corresponding single-run scores, which are multi-run DreamDIA deep learning score, multi-run spectral cosine similarity score, multi-run MS1 area score and multi-run MS2 area score. These multi-run scores can be an endorsement for well-aligned peak groups to be accepted by the statistical model. The weight of each single-run score in the linear combination is determined by the NC similarity of the run it belongs to the target run. This strategy helps to avoid low-quality peak groups from being picked and accepted just because their multi-run scores have been over-boosted by high-quality peak groups in other runs. To further minimize false information transfer between distant runs, we introduced an exponential weight decay function, *ϕ*:

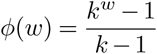

where *w* represents the single-run score weight, and *k* is a parameter named as weight decay coefficient. This function significantly penalizes lower weight values while having minimal impact on higher weight values. As a result, single-run scores from highly similar runs predominantly contribute to the multi-run score.

Ultimately, we obtain four single-run scores, four multi-run scores, and several additional simple scores, such as delta RT score, peptide length, and charge for each peak group, preparing them for subsequent statistical analysis. ***Statistical analysis.*** DreamDIAlignR constructs a semi-supervised learning model to distinguish between target peptides and decoys, utilizing either random forest or XGBoost algorithms. This approach mirrors the strategy employed by PyProphetRosenberger et al. (2017), with the key difference being the inclusion of multi-run scores in the analysis. The false discovery rate (FDR) is then estimated based on the distribution of discriminant scores between the target and decoy groups.

### Software information

DreamDIAlignR is integrated in the DreamDIAGao et al. (2021) software package (version 3.2.0). DreamDIAlignR was benchmarked with OpenSWATHHannes L Röst et al. (2014) (v2.7), DIAlignRGupta et al. (2019, 2023) (v2.6.0), DreamDIAGao et al. (2021) (v2.0.3) and DIA-NNDemichev et al. (2020) (v1.8).

## Data preprocessing

All the DIA raw data files were converted to open file format mzML using MSConvertChambers et al. (2012) (version: 3.0.23080). Format conversion was conducted twice: once with the “Peak Picking” filter to obtain centroided files for DIA-NN, DreamDIA and DreamDIAlignR, and once without the filter to produce profile files for OpenSWATH. Software parameters used for all the experiments are shown in Supplementary Note 2.

### Feasibility test on the *S. pyogenes* dataset

The *S. pyogenes* datasetHannes L Röst et al. (2014) was acquired on SCIEX TripleTOF 5600 instrument. It comprises a total of 16 runs, with 8 runs containing 10% human plasma as background and the remaining 8 runs lacking this component (Supplementary Figure S10a). A spectral library built in the original paperHannes L Röst et al. (2014); Röst et al. (2016); Gupta and Röst (2021) was used. We manually checked the peak boundary annotation file to discard ambiguously and falsely annotated peaks (see Supplementary Note 1). Eventually, 6870 peaks were retained in total including identification results from 434 peptides across 16 runs. Then we used the annotated peak locations to benchmark the peak identification performance of DreamDIAlignR. A correct identification is defined as an identified peak apex that falls into the range of the annotated peak boundaries.

### LFQbench test

The LFQbench HYE110 datasetNavarro et al. (2016), acquired on the SCIEX TripleTOF 6600 instrument, was chosen for benchmarking (Supplementary Figure S10b). To compare the quantification performance of different software tools across varying numbers of identified peptides, irrespective of hard FDR cutoffs, all tools were configured to output all results without applying any FDR filtering. Subsequently, the results were manually filtered using a range of FDR thresholds from 0.001 to 0.1. These filtered results were then input into the LFQbench software package to calculate quantification bias metrics, including 1 - species separation ability (SSA), median bias (MB), and dispersion (DISP). Species separation ability is defined as the area under the receiver operating characteristic (ROC) curve of a binary classifier between two species. Median bias is defined as the distance between the median of quantified Sample A-to-Sample B ratios and the ground truth ratio lines. Dispersion is defined as the standard deviation of the log-transformed ratios for each speciesNavarro et al. (2016). To illustrate overall quantification performance, we normalized the three quantification bias metrics using OpenSWATH as a baseline and defined total quantification bias as the mean of these three normalized metrics at 1% FDR. To account for the impact of varying FDR thresholds on the identification results of OpenSWATH with MBR, we executed DIAlignR six times after OpenSWATH analysis, each time specifying a different FDR threshold. We then defined a metric, Overall Performance Distance (OPD), to measure the overall identification and quantification performance improvement brought by the MBR approaches for different software tools. In this test, the overall performance of each software tool, with or without applying MBR algorithms at a specific FDR, is depicted by a 4-dimensional metric vector:

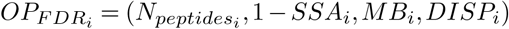

where the first item reflects identification performance and the other three reflect quantification performance. We first normalized all vectors for all software tools at all FDR thresholds to scale each item from 0 to 1. Then we calculated the Euclidean distance between each pair of metric points at the same FDR threshold, before and after the application of MBR for each software tool:

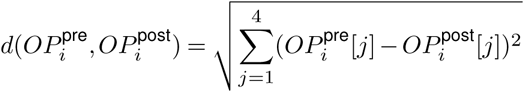

where *OP_i_*^pre^ and *OP_i_*^post^ are the normalized performance metric vectors before and after applying MBR at FDR threshold *i*, respectively. Then the OPD was calculated as the median of the distances across all the FDR thresholds for each software tool:

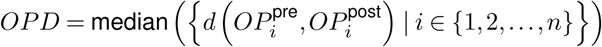

where *n* represents the number of FDR thresholds. To more reasonably represent performance enhancement across different FDR thresholds, the FDR thresholds were designed to be uniformly distributed in the logarithmic space. For comparison, OPDs were also calculated in the linear space (Supplementary Note 3). Specifically for OpenSWATH with MBR, the metrics were linearly interpolated based on the results of neighboring FDR thresholds if the results at specific FDR thresholds were absent.

### Two-sample, two-proteome test

To create a two-sample, two-proteome dataset, we selected 24 DIA runs from a previous published dataset which had been used to evaluate analysis reproducibility of large-scale DIA proteomics studiesPoulos et al. (2020) (referred to as TSTP dataset). The selected runs were technical replicates of two samples acquired on three different SCIEX TripleTOF 6600 mass spectrometers (#M2, #M4 and #M6) on two different days (Day14 and Day103). The 12 runs consisting of 50% prostate cancer tissue and 50% yeast cell lysates can be regarded as human-yeast mixed samples. Conversely, the remaining 12 runs with 50% ovarian cancer tissue and 50% prostate cancer tissue can be regarded as human-only samples (Supplementary Figure S10c). The human spectral library and the yeast spectral library were combined to get the mixed library for analysis. Peptide precursors in the library were filtered to have top 6 intensive fragment ions. All the software tools were configured to output identification results at 1% precursor FDR. Peptide precursors detected in less than one-third of the runs or with intensities below 100 were excluded from the benchmark analysis. The two-species FDR was calculated using the “combined” method as described by Wen et al.Wen et al. (2024), with the following formulas:

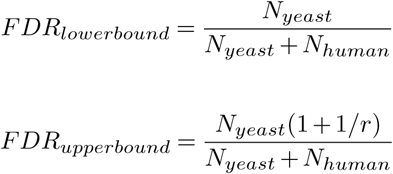

Here, *N_yeast_* and *N_human_* deonte the numbers of human and yeast peptides identified in all the runs without the yeast component. The parameter *r* represents the effective ratio of yeast peptides in the mixed spectral library, which is 0.267 in this experiment. The upper-bound FDR provides a more conservative estimation, though it risks overestimating the actual FDR. Conversely, the lower-bound FDR may overlook the differences in the likelihood of identifying human versus yeast peptidesWen et al. (2024). To provide a more comprehensive assessment, both upper-bound and lower-bound FDRs were presented.

### Heterogeneous dataset tests

To comprehensively benchmark the identification and quantification performance of DreamDIAlignR on highly heterogeneous dataset, we selected an additional 36 runs from the Procan cancer datasetPoulos et al. (2020) (referred to as Procan36 dataset). This dataset also includes replicates of two distinct samples. Specifically, 18 runs contain 6.25% ovarian cancer tissue, 50% prostate cancer tissue and 43.75% yeast cell lysates, while the other 18 runs contain 25%, 50% and 25% for the three portions respectively (Supplementary Figure S10d). Thus, the two sample sets allow us to establish ground truth species ratios (Ovary, 1:4; Prostate, 1:1; Yeast, 1.75:1) similar to those in the LFQbench dataset. The Procan36 dataset comprises DIA runs acquired on three different SCIEX TripleTOF 6600 mass spectrometers (#M1, #M3 and #M5) across two different days (Day14 and Day105). The diversity of cancer tissue introduces biological heterogeneity, while variations in experimental conditions result in technical heterogeneity. These factors collectively contribute to greater overall heterogeneity among runs compared to the LFQbench dataset. We computed quantification bias metrics, including median bias and dispersion, following the guidelines provided by the LFQbenchNavarro et al. (2016) software package. The total quantification bias and OPD were calculated in a manner similar to the LFQbench test, but without including species separation ability to avoid introducing bias from different metric calculation methods for the benchmark. Additionally, we defined ovary-specific peptides as those identified by DIA-NN (at a 1% FDR) in at least 8 out of 12 Ovary 50% runs but in at most 1 out of 12 Ovary 0% runs in the TSTP dataset. For data matrix completeness benchmarking, We filtered the quantification matrices yielded by all the software tools to include only analytes that had been identified in at least three runs and defined data completeness as the number of validly quantified entries divided by the total number of entries.

Furthermore, we selected a dataset of 494 runs from the Procan dataset to evaluate performance on a large-scale data (referred to as Procan494 dataset). This dataset includes replicates of four distinct samples, each with varying organism/species proteome ratios (Supplementary Figure S10e). The data were acquired using 6 different SCIEX TripleTOF 6600 mass spectrometers (#M1, #M2, #M3, #M4, #M5, and #M6) across 7 different days (Day7, Day14, Day21, Day28, Day56, Day84, and Day107). To assess quantification performance, we calculated the median of the *R*^2^ values from the calibration curves for both ovarian peptides and yeast peptides. For identification performance, we defined a valid peptide as one identified in at least ten runs within each sample and then calculated the total number of valid peptides identified.

### Differential expression analysis and over-representation analysis

Differential expression analysis was performed on the TSTP dataset with a widely-accepted workflowPeng et al. (2024) as follows. Protein quantification matrices were generated by summing the peak areas of the top three most intense peptide precursors for each protein across DreamDIAlignR, DIA-NN with MBR, and OpenSWATH with MBR, all at 1% precursor FDR. Human peptides with intensities lower than 100, as well as all yeast peptides, were excluded. Additionally, peptide precursors identified in fewer than one-third of the runs were discarded. The remaining quantification matrices were imputed using k-nearest neighbors (KNN) algorithm with three neighbors. Following imputation, the matrices were quantile normalized, log-transformed, and then analyzed with limmaRitchie et al. (2015) for protein signature identification and statistical analysis.

Lastly, we utilized the ClusterProfilerYu et al. (2012); Wu et al. (2021); Xu et al. (2024) R package to perform over-representation analysis for the differentially expressed proteins using two different databases: Disease Ontology (DO)Schriml et al. (2022) and ProteomicsDBSamaras et al. (2020); Chen et al. (2013). For the DO analysis, we focused on ovarian cancer-related DO terms to facilitate comparison across different software tools. In the ProteomicsDB enrichment analysis, we selected the top five enriched gene sets with adjusted p-values below 0.1 as benchmarks for evaluation.

## Data availability

All the datasets used in this work are published and publicly available. The *S. pyogenes* dataset was deposited to PeptideAtlas with accession code PASS00788. The LFQbench dataset and the Procan dataset were deposited to ProteomeXchange ConsortiumMa et al. (2019); Perez-Riverol et al. (2019) with dataset identifiers PXD002952 and PXD015912, respectively. The code and data used to generate the figures have been uploaded to Zenodo (10.5281/zenodo.14531668).

## Code availability

DreamDIAlignR is fully open-source and available at https://github.com/xmuyulab/DreamDIA.

## Acknowledgement

This work was supported by Canadian Institutes of Health Research (Grant No. 507496) and China Scholarship Council (Grant No. 202206310091).

## Author contributions

M.G., H.R. and R.Y. designed the study. M.G., S.G. and W.Y. implemented the algorithms. M.G. analyzed the data.

M.G. wrote the first manuscript. H.R. and R.Y. supervised the study.

## Supplementary Notes

**Supplementary Note 1:**
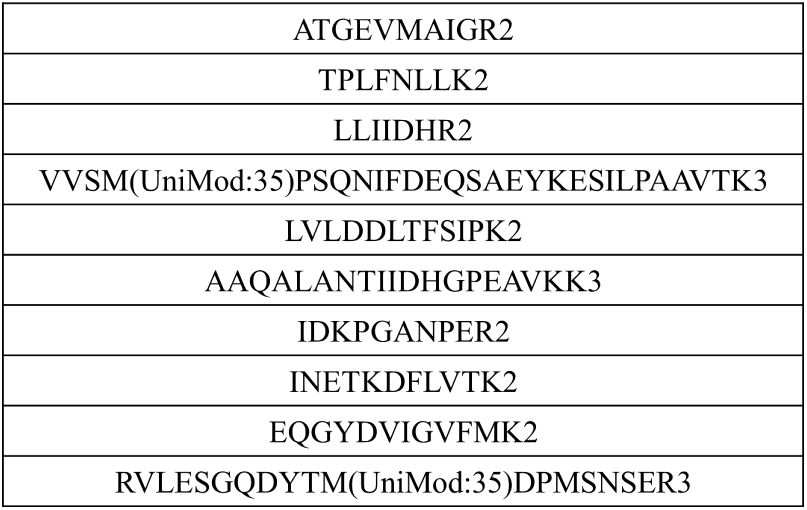
Excluded Peptide Annotations in S.pyogenes Dataset. We double-checked the manually annotated peak boundaries of the 400 peptides in S.pyogenes dataset by previously published work. We found that some of the annotated peak boundaries were confusing or wrong. Thus, we chose to exclude the following peptide annotations in our benchmark experiments.

**Supplementary Note 2:**
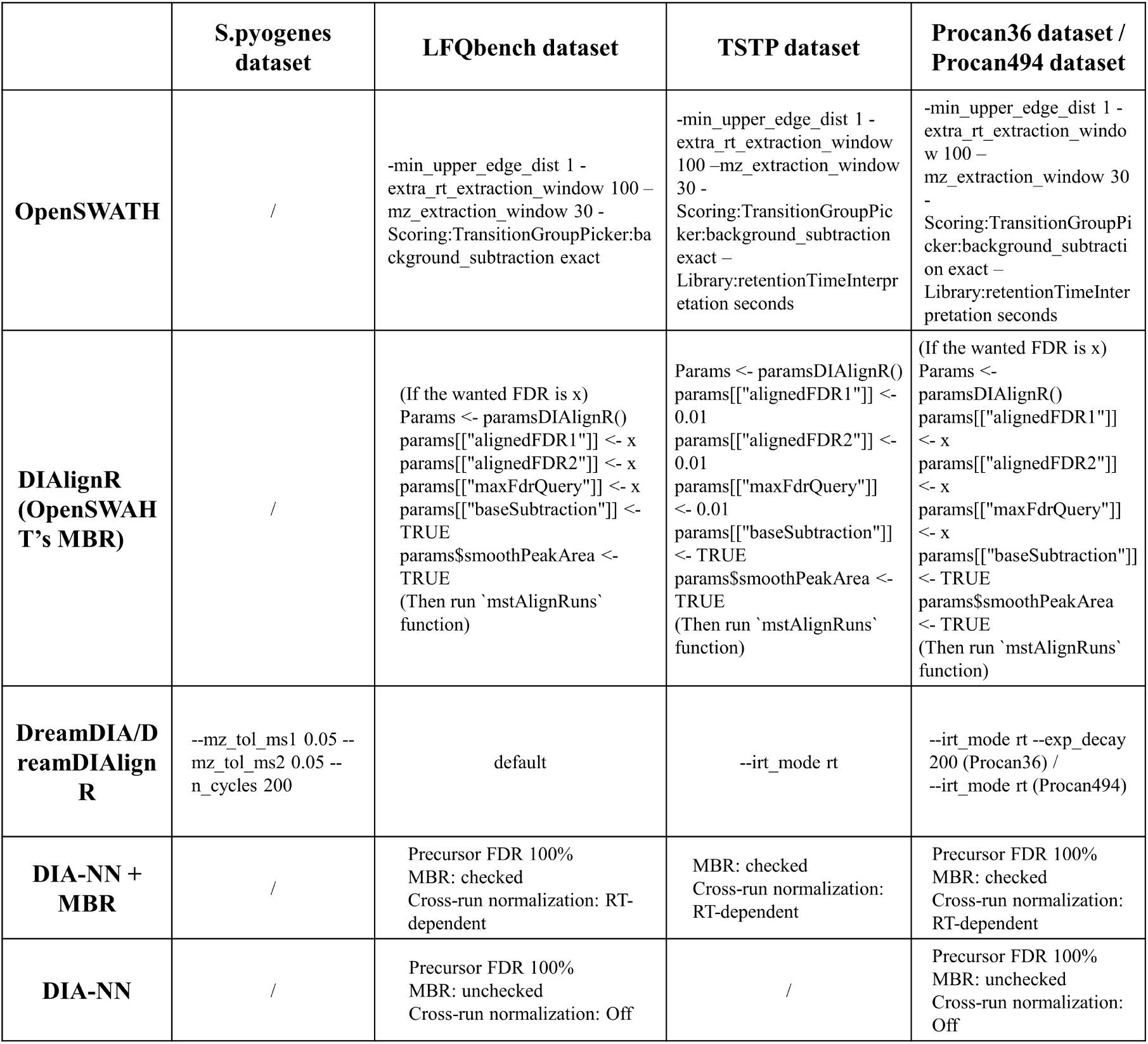
Software Parameters

**Supplementary Note 3:**
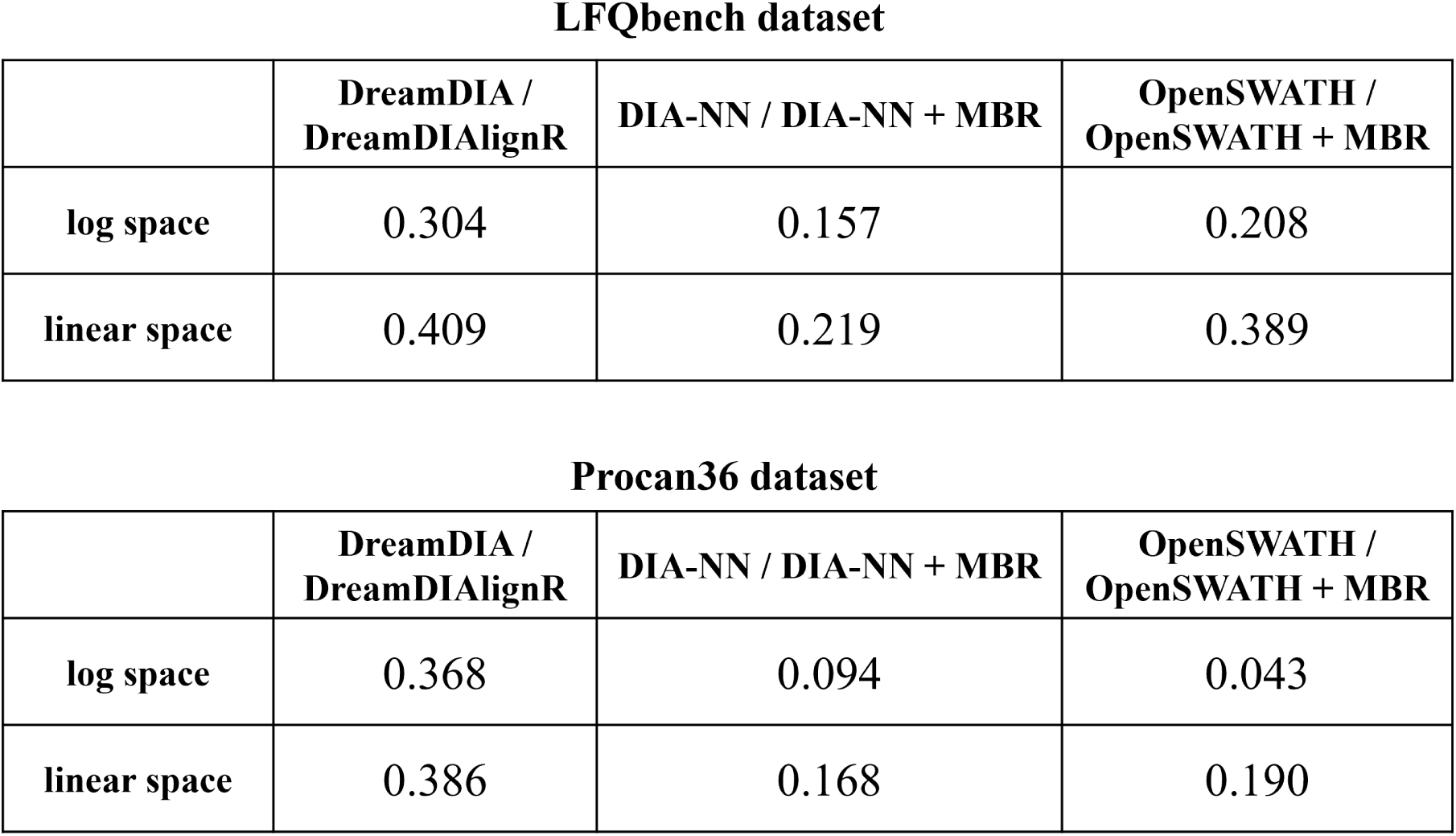
Overall Performance Distance (OPD) benchmark results

## Supplementary Figures

**Figure S1.**
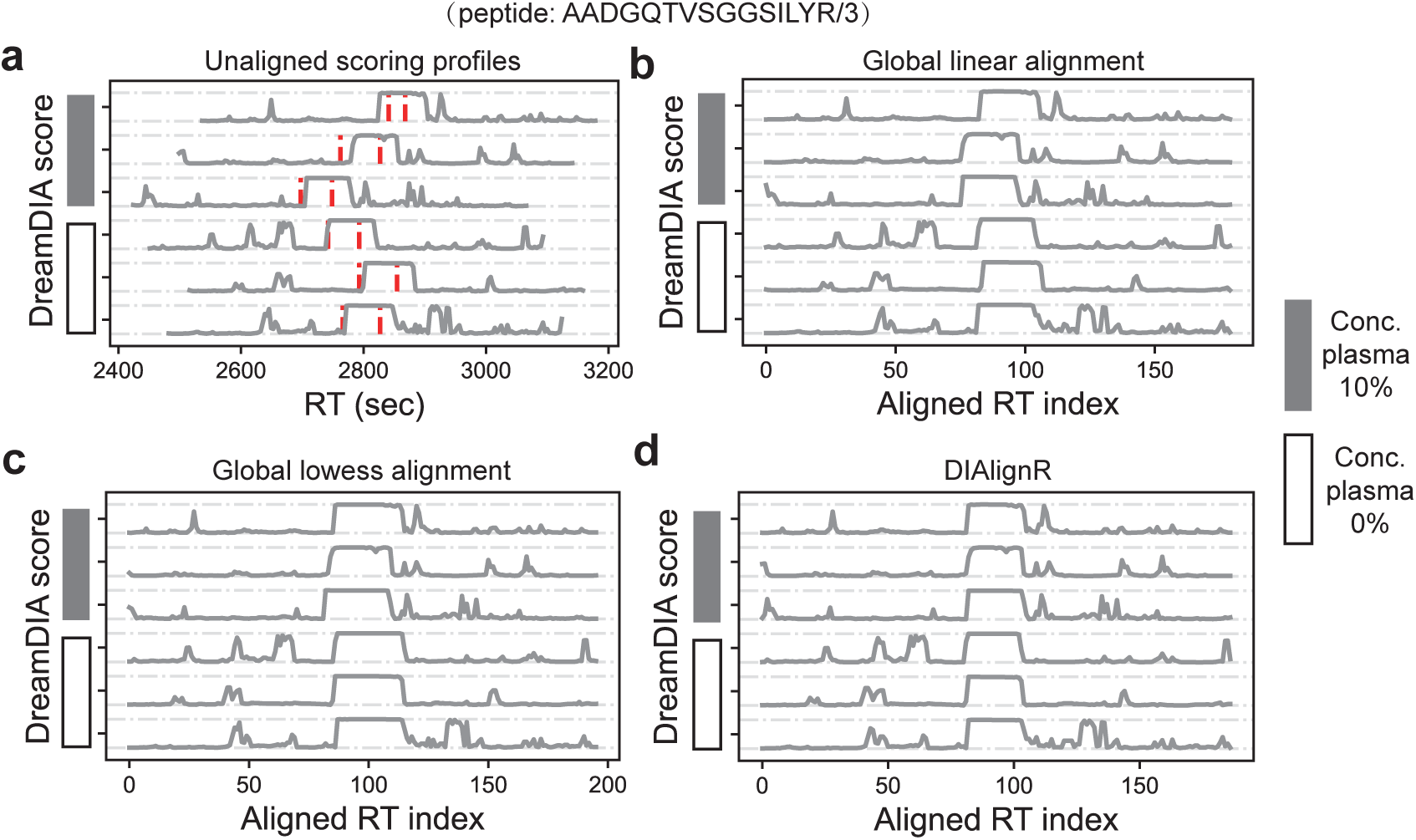
Scoring profile sychronization performance of an example peptide AADGQTVSGGSILYR/3 in Streptococcus Pyogenes dataset using different signal alignment methods. **a.** Unaligned scoring profiles. **b.** Aligned scoring profiles using linear global alignment. **c.** Aligned scoring profiles using lowess global alignment. **d.** Aligned scoring profiles using DIAlignR. Red dashed lines denote manually annotated peak boundaries. Only results of 6 out of 16 runs are shown due to the limit of figure space.

**Figure S2.**
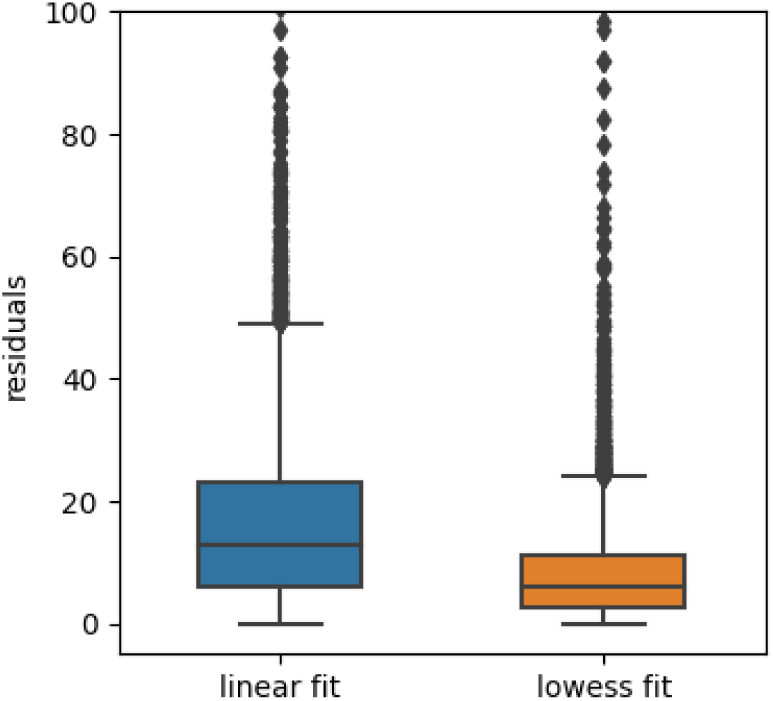
Residuals of linear or lowess global alignment methods on S.pyogenes dataset. A linear or lowess fit was built between the RT values of anchor peptides from each run pair in the global minimum spaning tree. Residuals of all the anchor peptides in all the fit models were plotted. Residuals higher than 100 were discarded. Boxplot elements: center line, median; boxes, interquartile range; whiskers, 1.5x interquartile range; points, outliers.

**Figure S3.**
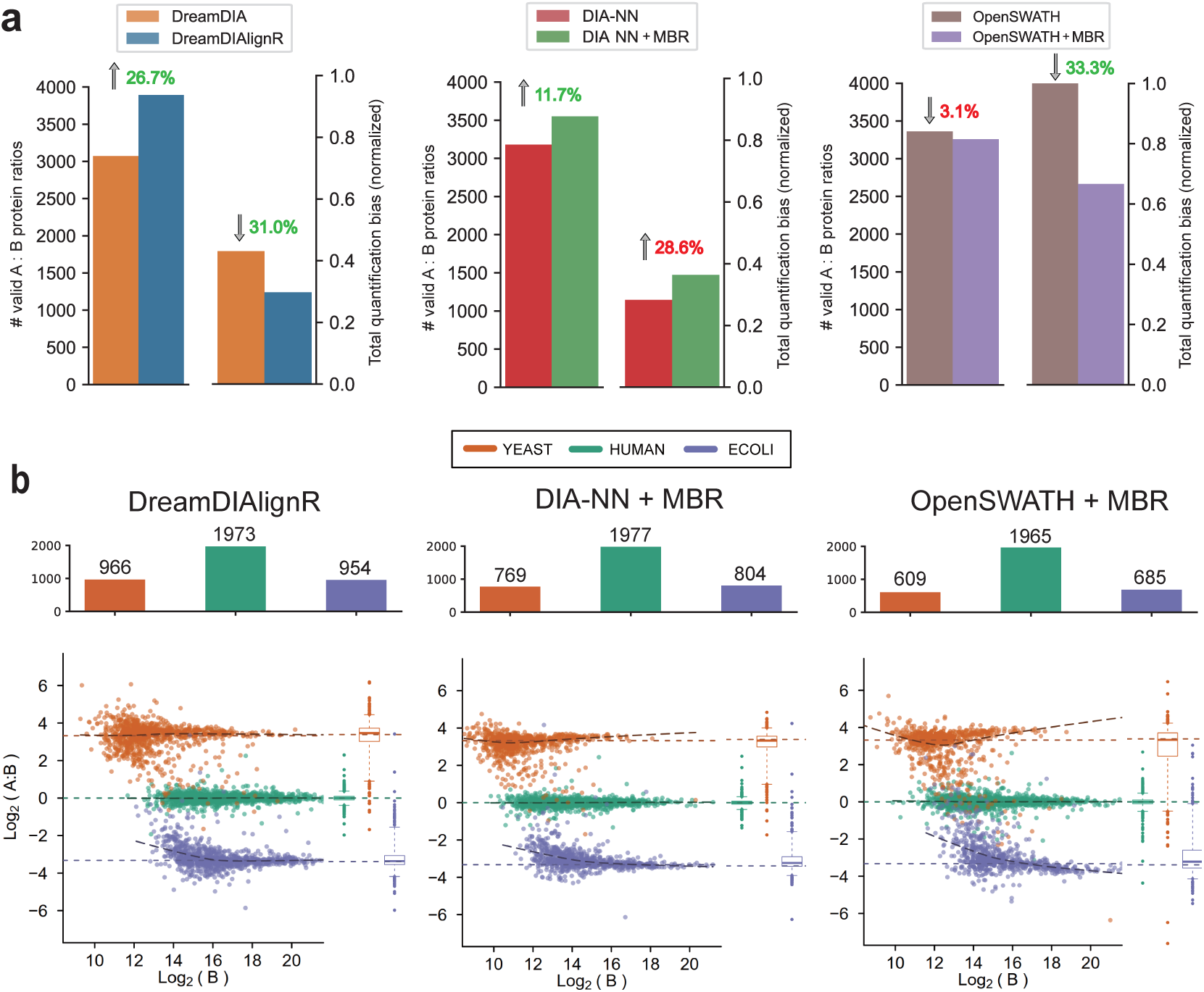
Identification and quantification performance benchmark on LFQbench dataset (protein level). **a.** The number of valid protein ratios identified and the total quantification bias of various software tools before and after applying multi-run alignment at 1% precursor FDR. Total quantification bias is calculated as the mean of the three normalized quantification performance metrics given by LFQbench software suite including 1 - species separation ability, median bias and dispersion (See Methods). **b.** Protein-level LFQbench benchmark results of OpenSWATH + Match-Between-Runs (MBR), DIA-NN + MBR and DreamDIAlignR at 1% precursor FDR. Colored dashed lines denote log-transformed ground truth Sample A to Sample B ratios (Human, 1:1; Yeast, 10:1; E.coli, 1:10). The numbers of valid protein ratios identified for different species are shown in the bar charts. Boxplot elements: center line, median; boxes, interquartile range; whiskers, 1.5x interquartile range; points, outliers.

**Figure S4.**
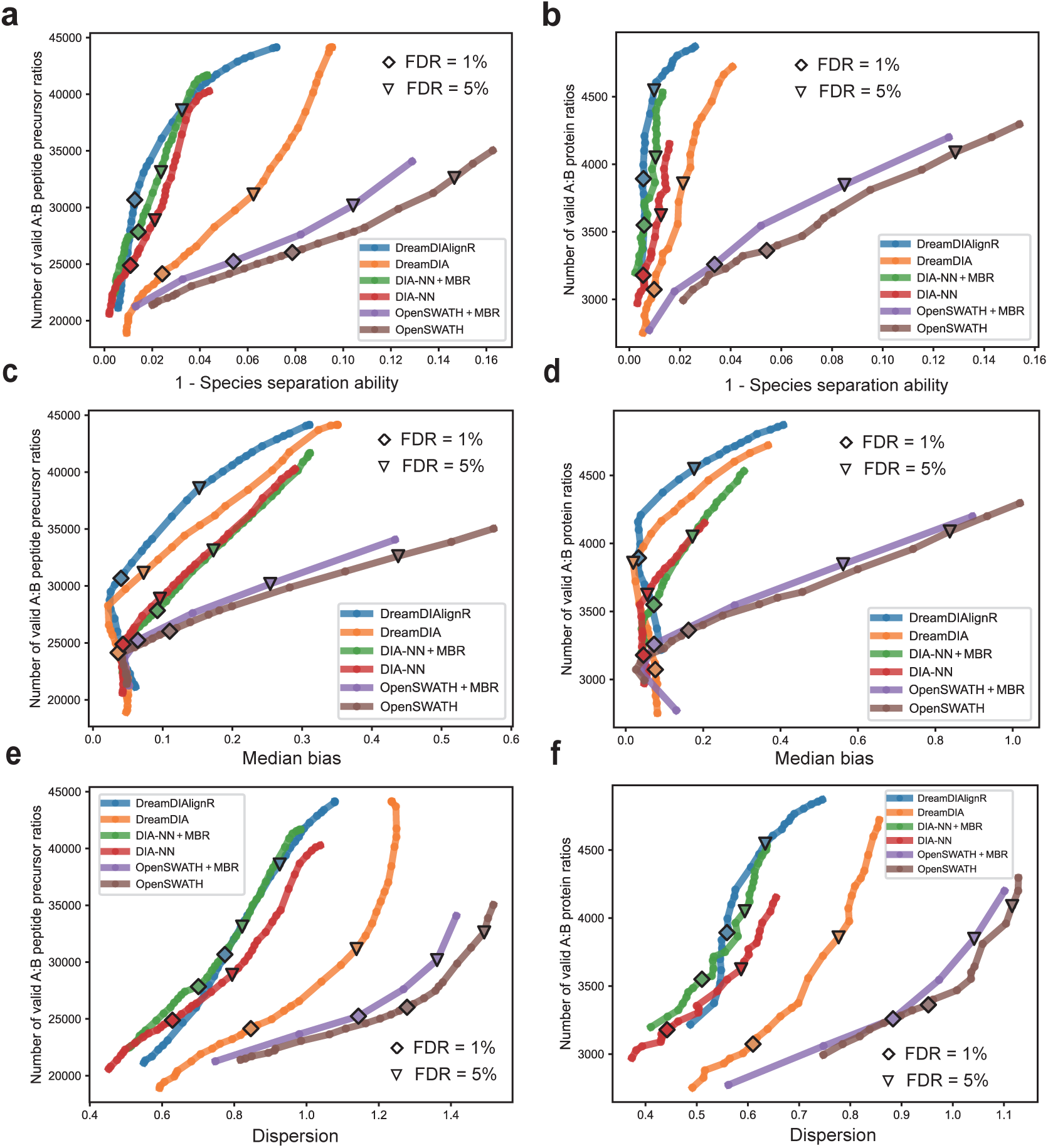
Comprehensive identification and quantification performance benchmark on LFQbench dataset. The number of valid Sample A to Sample B peptide precursor ratios (**a, c, e**) and protein ratios (**b, d, f**) identified in total are plotted against the corresponding three quantification bias metrics including 1 - species separation ability (**a, b**), median bias (**c, d**) and dispersion (**e, f**) using a series of FDR thresholds. A peptide can only be designated as having a valid ratio if it has been identified in at least one Sample A run and one Sample B run. “♢” and “▽” denote the results at 1% and 5% peptide precursor FDR, respectively.

**Figure S5.**
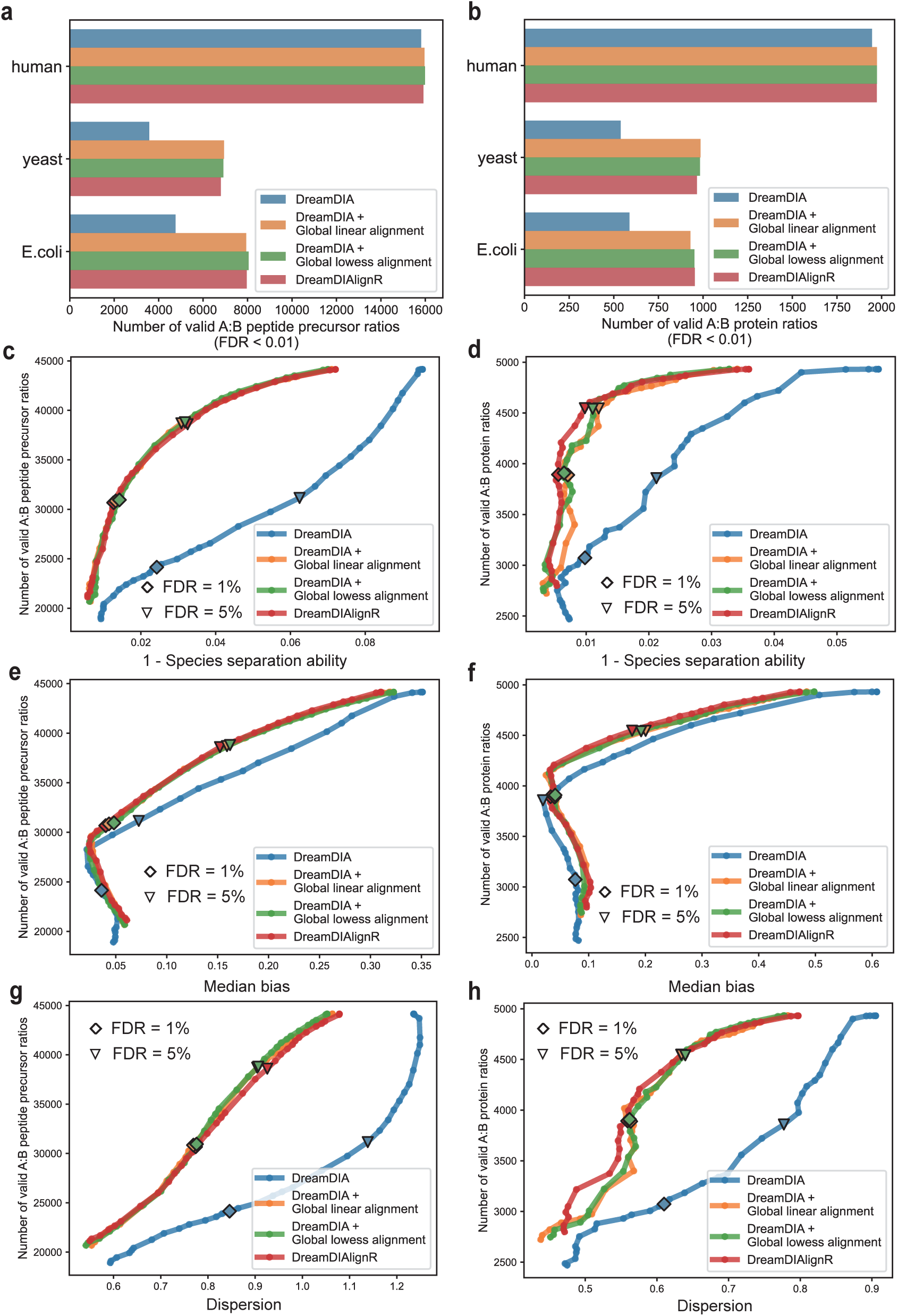
Identification and quantification performance benchmark of DreamDIAlignR using different signal alignment methods on LFQbench dataset. a, **b.** The number of peptide precursors (**a**) and proteins (**b**) identified at 1% precursor FDR. **c-h.** The number of valid Sample A to Sample B peptide precursor ratios (**c, e, g**) and protein ratios (**d, f, h**) identified in total are plotted against the corresponding three quantification bias metrics including 1 - species separation ability, median bias and dispersion using a series of FDR thresholds.

**Figure S6.**
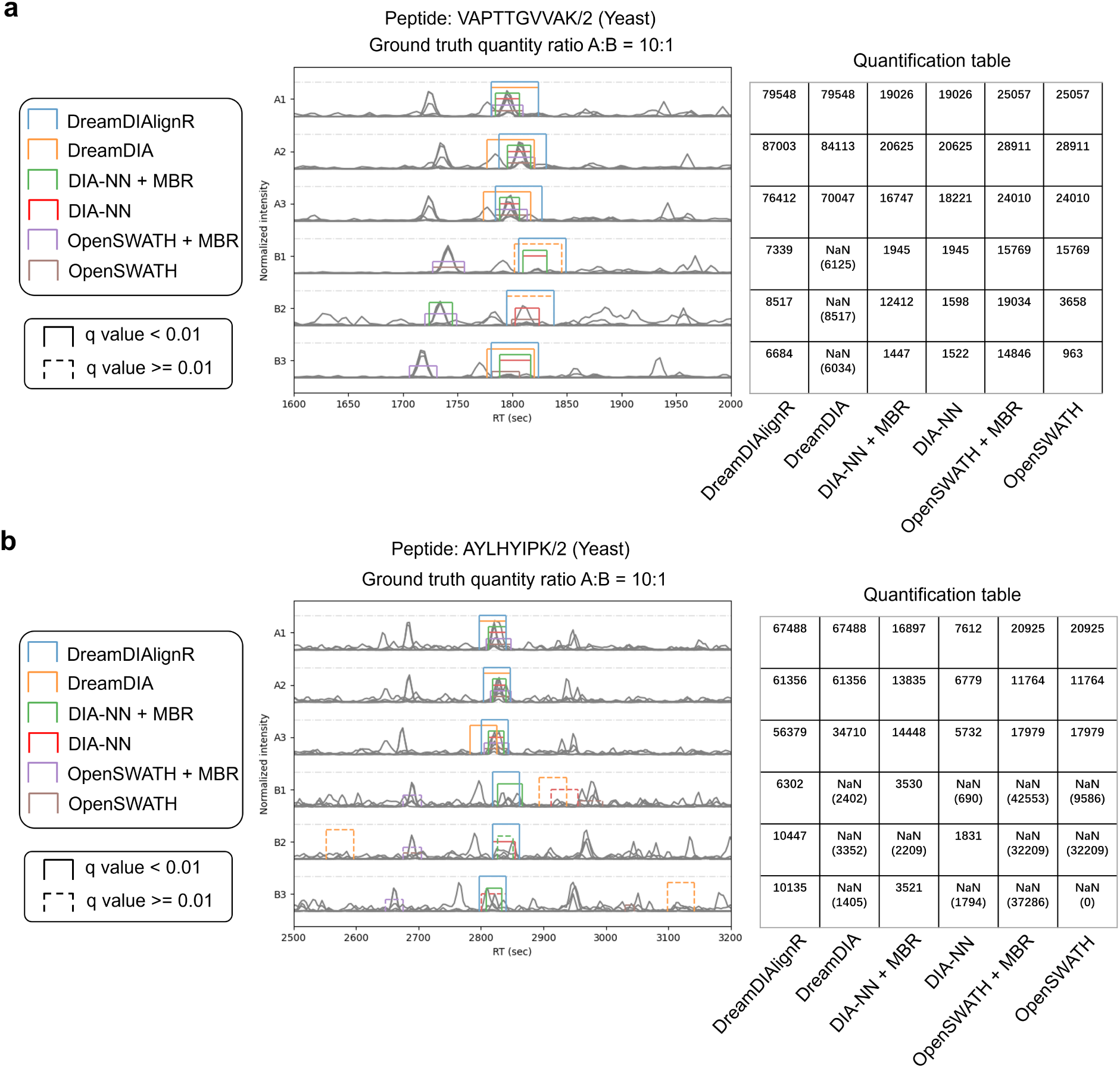
Example peak groups identified using different software tools on LFQbench dataset. **a.** Peak groups identified for peptide VAPTTGVVAK/2. **b.** Peak groups identified for peptide AYLHYIPK/2. Solid lines and dashed lines stand for peak groups with *q* values lower or higher than 1% respectively. Peptide quantities of each run given by different software are attached beside the corresponding chromatograms.Quantities corresponding to peak groups with *q*-values above 1% are displayed in parentheses, indicating their invalidity for reliable quantification. OpenSWATH requires a single peak group from all runs to be picked as the reference peak group and subsequently map this reference to neighboring runs. However, if the reference peak group is incorrectly picked (Run B1 in (**a**) and Run B2 in (**b**)), it reverberates across all related runs, potentially compromising the entire set of peak groups. DreamDIAlignR takes a different approach. It aggregates peak groups from all runs, leveraging the averaged cross-run scoring profile. By reducing reliance on a single reference run, DreamDIAlignR achieves heightened consistency and cross-run quantification accuracy. Moreover, the quantification matrix generated by DreamDIAlignR is also more complete with fewer missing values since the well-aligned high-quality peak groups are assigned more reasonable scores to be taken into consideration for quantification.

**Figure S7.**
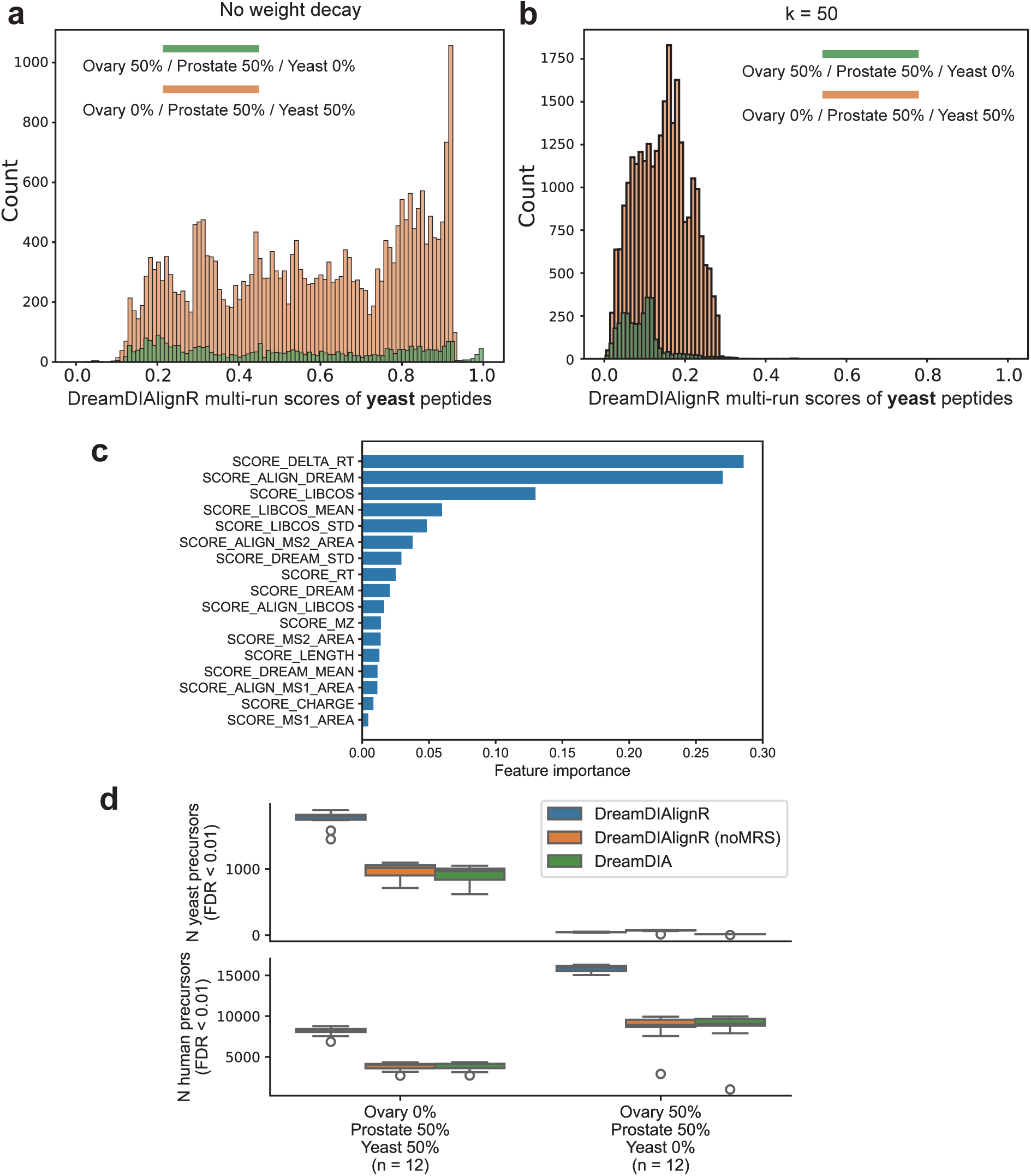
Optimization of the weight decay parameter for multi-run score (MRS) calculation on TSTP dataset. a, **b.** Distributions of the multi-run scores of yeast peptides in the two sample sets without (**a**) or with (**b**) weight decay. **c.** Feature importance of the statistical model when analyzing the TSTP dataset. The default XGBoost model was used. Scores with the prefix “SCORE_ALIGN” represent the multi-run scores calculated in DreamDIAlignR. **d.** The numbers of human and yeast peptide precursors identified by DreamDIA and DreamDIAlignR with and without the multi-run scores used by the discriminative model. Boxplot elements: center line, median; boxes, interquartile range; whiskers, 1.5x interquartile range; points, outliers.

**Figure S8.**
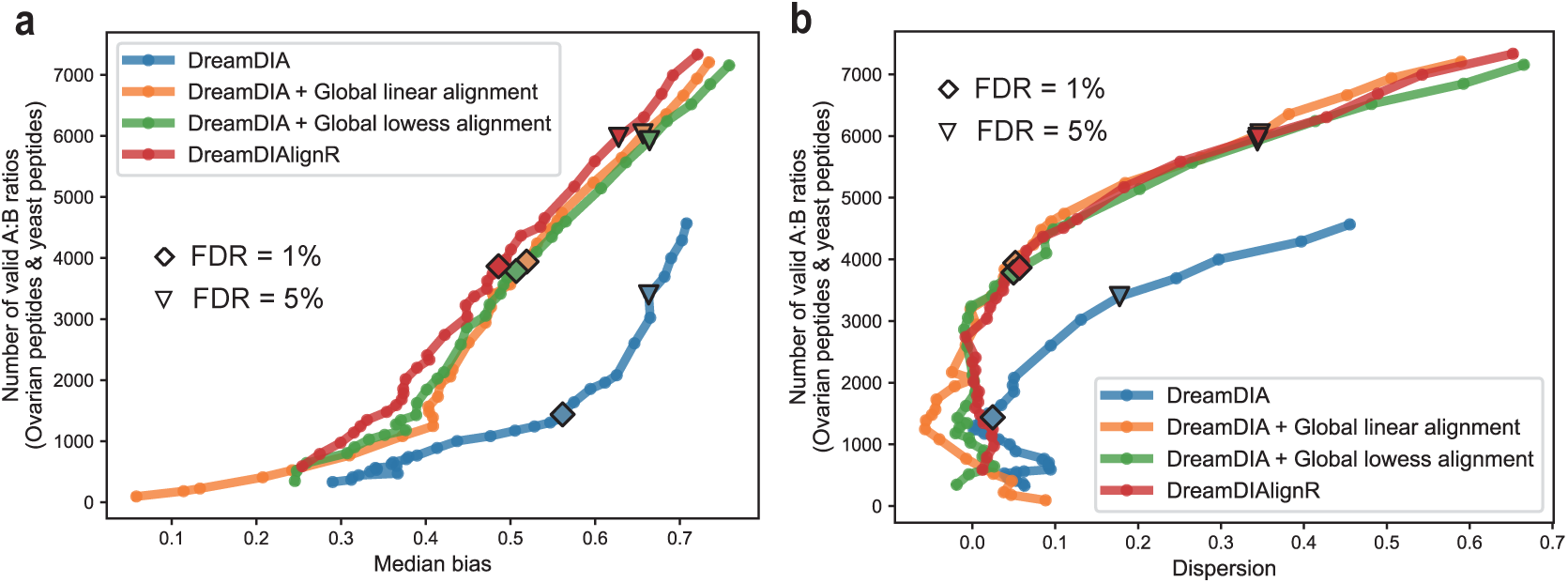
Performance of different signal alignment methods when analyzing highly heterogeneous dataset. The number of valid yeast and ovarian Sample A to Sample B peptide precursor ratios is plotted against the corresponding quantification bias metrics, median bias (**a**) and dispersion (**b**) using a series of FDR thresholds. A peptide is named as a valid ratio only if it has been identified in at least 3 Sample A runs and 3 Sample B runs. The quantification bias metrics were calculated as suggested by LFQbench software package. “♢” and “▽” mean the results at 1% and 5% peptide precursor FDR respectively.

**Figure S9.**
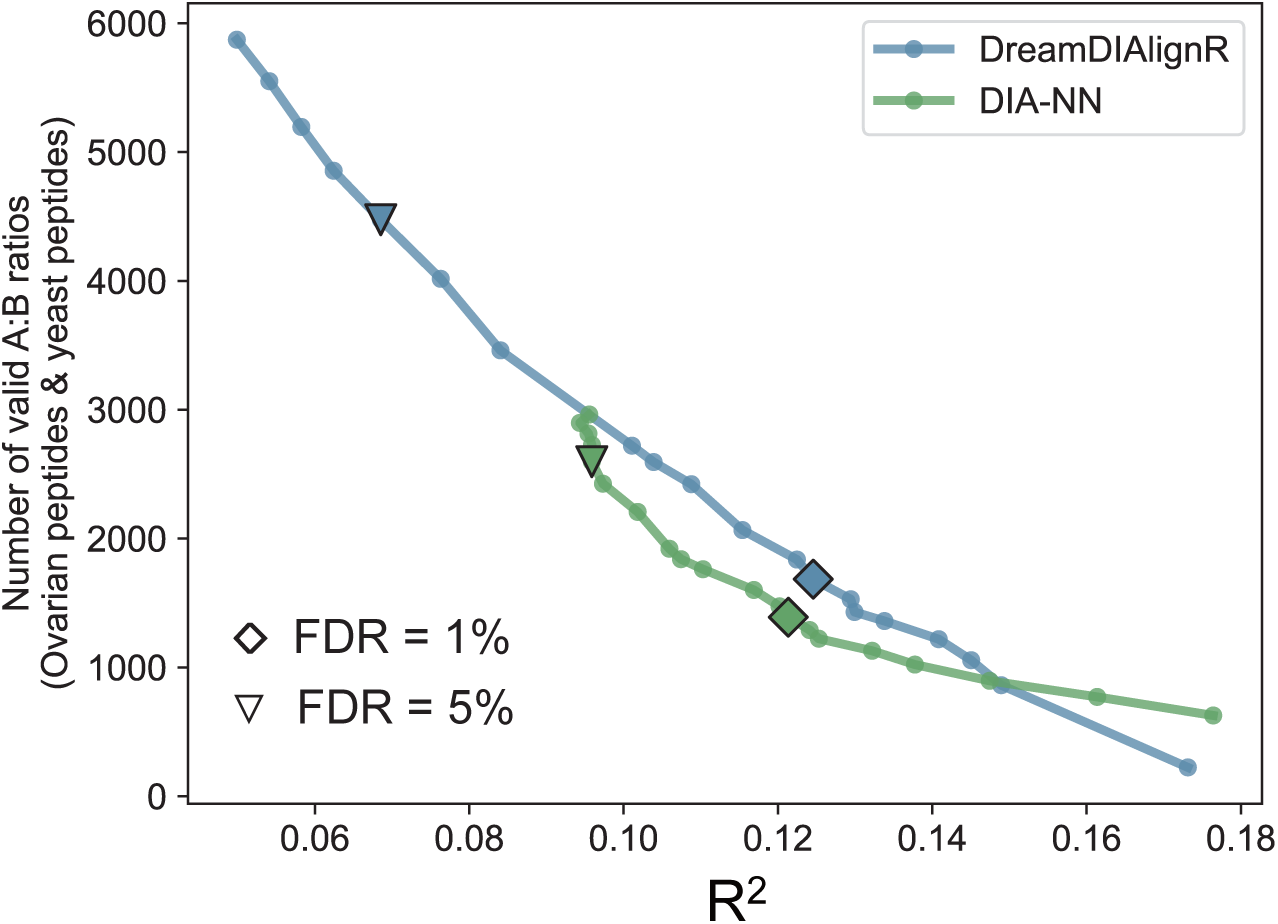
Performance of DreamDIAlignR and DIA-NN on Procan494 dataset. The number of valid peptide precursor ratios is plotted against the corresponding quantification metric, the median of *R*^2^, using a series of FDR thresholds. A valid ratio is defined as a peptide that has been identified in at least 10 runs for each sample. “♢” and “▽” denote the results at 1% and 5% peptide precursor FDR, respectively.

**Figure S10.**
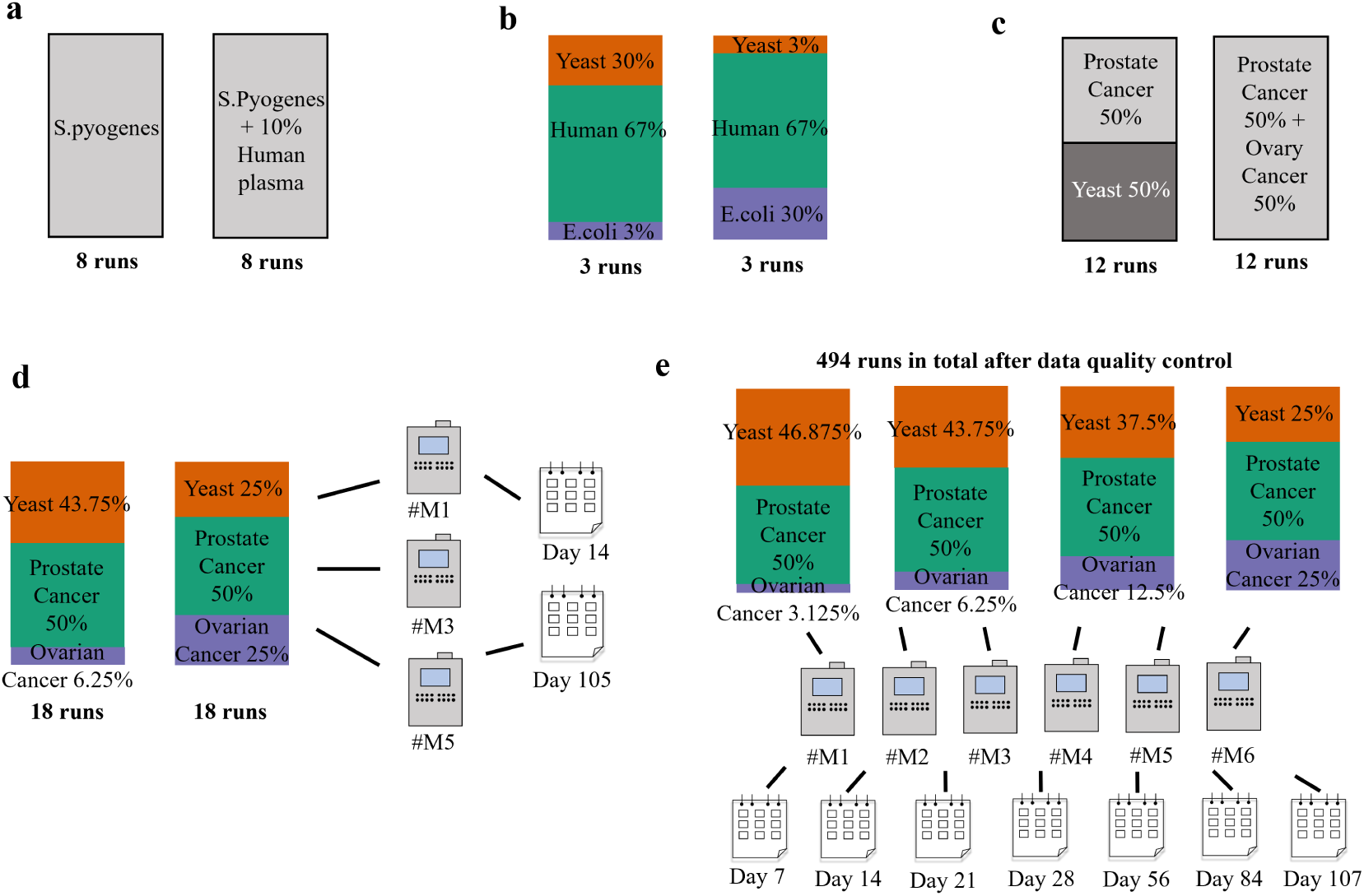
Schematic overview of the datasets analyzed in this study. (**a**) S.pyogenes dataset. (**b**) LFQbench HYE110 dataset. (**c**) TSTP dataset. (**d**) Procan36 dataset. (**e**) Procan494 dataset.

**Figure S11.**
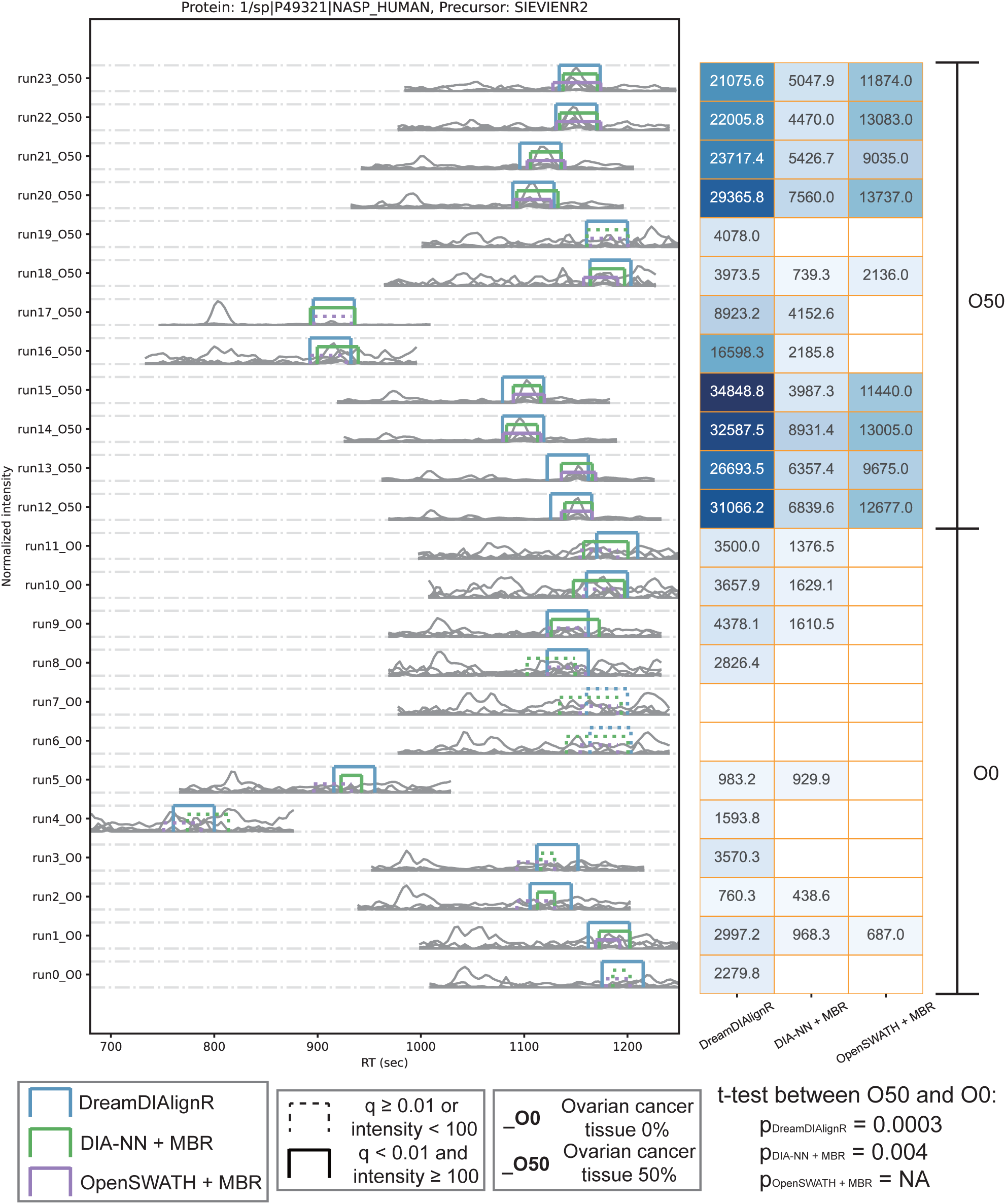
Chromatographic peaks of the precursor SIEVIENR2 from the protein NASP identified by three different software tools and their corresponding quantifications across 24 runs in the TSTP dataset. Colored boxes represent the peak boundaries assigned by each software tool. Solid lines indicate valid identifications, defined as having a q-value below 1% and intensity above 100, whereas dashed lines denote invalid identifications. Adjacent run pairs (e.g., even and odd runs such as Run0 and Run1, Run2 and Run3, Run4 and Run5) are technical replicates acquired under identical experimental conditions. T-tests were conducted between the precursor intensities of the “O50” and the “O0” groups without intensity imputation or normalization, with intensities in the “O0” group multiplied by 2 for fair comparison.

**Figure S12.**
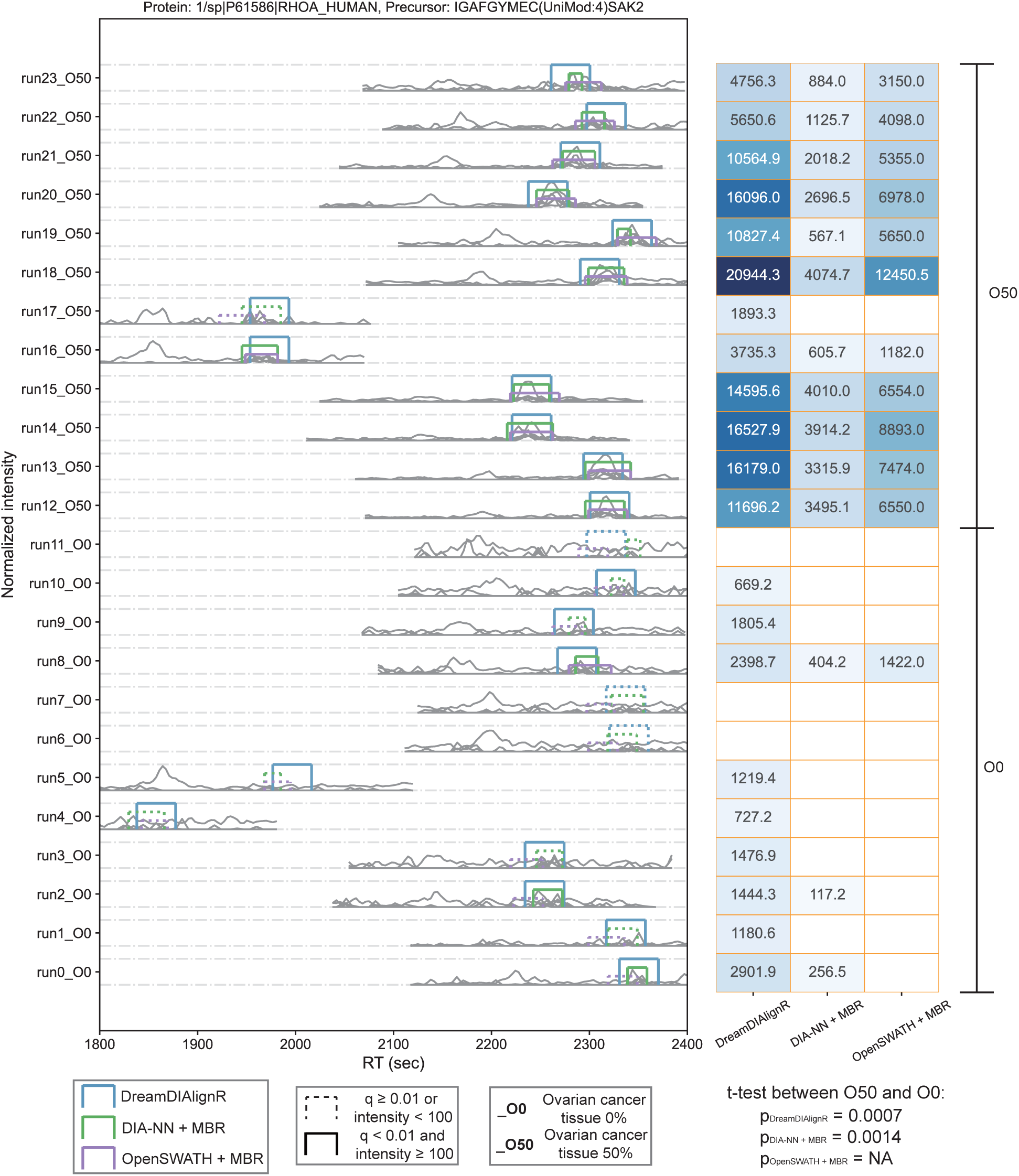
Chromatographic peaks of the precursor IGAFGYMEC(UniMod:4)SAK2 from the protein RHOA identified by three different software tools and their corresponding quantifications across 24 runs in the TSTP dataset. Colored boxes represent the peak boundaries assigned by each software tool. Solid lines indicate valid identifications, defined as having a q-value below 1% and intensity above 100, whereas dashed lines denote invalid identifications. Adjacent run pairs (e.g., even and odd runs such as Run0 and Run1, Run2 and Run3, Run4 and Run5) are technical replicates acquired under identical experimental conditions. T-tests were conducted between the precursor intensities of the “O50” and the “O0” groups without intensity imputation or normalization, with intensities in the “O0” group multiplied by 2 for fair comparison.

